# Comparative Analysis of Polysaccharide and Cell Wall Structure in *Aspergillus nidulans* and *Aspergillus fumigatus* by Solid-State NMR

**DOI:** 10.1101/2024.08.13.607833

**Authors:** Isha Gautam, Jayasubba Reddy Yarava, Yifan Xu, Reina Li, Faith J. Scott, Frederic Mentink-Vigier, Michelle Momany, Jean-Paul Latgé, Tuo Wang

## Abstract

Invasive aspergillosis poses a significant threat to immunocompromised patients, leading to high mortality rates associated with these infections. Targeting the biosynthesis of cell wall carbohydrates is a promising strategy for antifungal drug development and will be advanced by a molecular-level understanding of the native structures of polysaccharides within their cellular context. Solid-state NMR spectroscopy has recently provided detailed insights into the cell wall organization of *Aspergillus fumigatus*, but genetic and biochemical evidence highlights species-specific differences among *Aspergillus* species. In this study, we employed a combination of ^13^C, ^15^N, and ^1^H-detection solid-state NMR, supplemented by Dynamic Nuclear Polarization (DNP), to compare the structural organization of cell wall polymers and their assembly in the cell walls of *A. fumigatus* and *A. nidulans*, both of which are key model organisms and human pathogens. The two species exhibited a similar rigid core architecture, consisting of chitin, α-glucan, and β-glucan, which contributed to comparable cell wall properties, including polymer dynamics, water retention, and supramolecular organization. However, differences were observed in the chitin, galactosaminogalactan, protein, and lipid content, as well as in the dynamics of galactomannan and the structure of the glucan matrix.

## Introduction

Among the vast diversity of fungi, the genus *Aspergillus* stands out as one of the most abundant and ubiquitous saprophytes, comprising 339 known species in the Trichocomaceae family (Houbraken et al., 2020). Fewer than 20 of these species, including *A. fumigatus*, *A. flavus*, *A. terreus*, *A. niger*, *A. nidulans*, *A. oryzae*, and *A. parasiticus*, are known human pathogens. *A. fumigatus* is the most prevalent, responsible for approximately 90% of all aspergillosis cases, particularly affecting individuals with compromised immune systems or respiratory conditions (Denning, 1998; Latgé, 1999; Latgé & Chamilos, 2019). *A. nidulans* is recognized not only for its pathogenicity, especially in immunocompromised patients with chronic granulomatous disease (CGD) (Åhlin et al., 1995; Dotis & Roilides, 2004; Henriet et al., 2012), but also as a model genetic system (Borgia & Dodge, 1992; Galagan et al., 2005; Osmani & Mirabito, 2004). This species has been instrumental in studying cell cycle control, DNA repair, mutation recombination, cytoskeletal function, mitochondrial DNA structure, and human genetic diseases.

The fungal cell wall, a dynamic organelle composed of complex carbohydrates and proteins, is essential for maintaining cellular integrity, morphology, and interactions with the environment (Gow et al., 2017; Gow & Lenardon, 2023; Latgé & Chamilos, 2019). The cell wall also serves as important targets for the development of antifungal drugs, with the success of echinocandins, a class of antifungals targeting the biosynthesis of β-1,3-glucan for function (Bowman et al., 2002; Perlin, 2011). Given the species-specific growth characteristics and pathogenic determinants of *Aspergillus* species, understanding the structural and biosynthetic differences between the cell walls of key *Aspergillus* species is essential for guiding the development of cell wall-targeting antifungal drugs that are effective across all *Aspergillus* species. While the cell wall of *A. nidulans* shares some similarities with that of *A. fumigatus*, it also exhibits unique features that reflect its specific evolutionary adaptations and ecological niche (Galagan et al., 2005).

Previous chemical analyses of *A. nidulans* cell walls have identified a predominant presence of glucose and acetylglucosamine, along with mannose, galactose, galactosamine, proteins, and lipids (Fontaine et al., 2011). Comparative analyses of *Aspergillus* species have consistently shown a similar composition, where each fungus displaying approximately 40% α-glucan and 40% β-glucan (Gastebois et al., 2009; Guest & Momany, 2000). There are, however, discrepancies in the cell wall composition of these two *Aspergillus* species. The relative proportions of galactosamine (GalN) and N-acetyl galactosamine (GalNAc) between the cell walls of *A. nidulans* and *A. fumigatus*, with the latter containing a low level of GalNAc that is absent in *A. nidulans* (Guest & Momany, 2000; M. J. Lee et al., 2015). Furthermore, the overexpression of the α-glucan synthetase gene *agsB*, or the deletion of *UgeA* (UDP-glucose-4-epimerase) and *UgmA* (UDP-galactopyranose mutase) in *A. nidulans* has been linked to increased hyphal adhesion to hydrophobic surfaces and enhanced fungal virulence (He et al., 2013; He et al., 2018; Paul et al., 2011). These results suggest that the cell wall composition may differ between these two *Aspergillus* species.

Solid-state NMR spectroscopy is a non-destructive, high-resolution technique (Reif et al., 2021) that enables the examination of intact cells without the need for chemical perturbation or extraction (Ghassemi et al., 2022; Latgé & Wang, 2022). This technique has recently proven valuable in complementing conventional chemical assays, providing a comprehensive view of cell wall architecture in *Cryptococcus* species (Chatterjee et al., 2015; Chatterjee et al., 2018; Chrissian et al., 2020), *Schizophyllum commune* (Ehren et al., 2020; Kleiburg et al., 2023; Safeer et al., 2023), *Candida albicans* (Fernando et al., 2022), *Neurospora crassa* (Delcourte et al., 2024), as well as *A. fumigatus* mycelia and conidia (Chakraborty et al., 2021; Lamon et al., 2023). Since the cell wall composition of *A. nidulans* has not been thoroughly studied, we conducted a comparative analysis of *A. fumigatus* and *A. nidulans* using ^13^C and ^15^N solid-state NMR techniques, supplemented by proton detection via fast magic-angle spinning (MAS) (Marchand et al., 2022), sensitivity-enhancing dynamic nuclear polarization (DNP) (Biedenbander et al., 2022; Chow et al., 2022; D. Lee et al., 2015; Ni et al., 2013), and transmission electron microscopy (TEM). We focus on the mycelial cell walls, which are the infective propagules, to pinpoint specific differences that could lead to the identification of unique virulence determinants in these two species.

## Materials and Methods

### Preparation of ^13^C,^15^N-labeled fungal material

Two strains of *A. fumigatus*, Af293 and CEA17Δ*akuB*^KU80^ (Chakraborty et al., 2021) were cultured in ^13^C, ^15^N-enriched media containing 1% ^13^C-glucose and 0.6% ^15^N-NaNO_3_ as carbon and nitrogen sources, respectively. The media was supplemented with 1 mL/L of trace-element solution (0.04% Na_2_B_4_O_7_·10H_2_O, 5 mM FeCl_3_) and 0.2 M HCl for preventing oxidation. Additionally, a solution containing 0.05% KCl, 0.08% MgSO_4_·7H_2_O, and 0.11% KH_2_PO_4_ was added. The pH was adjusted to 6.5. The *A. fumigatus* strains were cultured in 100 mL liquid media in 250 mL Erlenmeyer flasks at 30 °C and 210 rpm. Fungal material was collected by washing with nanopure water and centrifuging at 7000 rpm (13700 × g). Similarly, *A. nidulans* (strain A28) was grown in minimal media (0.5% peptone, 1% complete supplement, 0.5% vitamin supplement), with the pH adjusted to 6.6. The composition of the medium is provided in **Supplementary Table 1**. Af293 was also cultured under the same conditions as A28. Cultures were incubated at 31°C for three days and washed five times with nanopure water to remove excess small molecules and reduce ion concentration. For each sample, 35-45 mg of whole-cell material was packed into a 3.2 mm MAS rotor for solid-state NMR characterization.

### Imaging of cell wall thickness and morphology

Three *Aspergillus* samples (A28, Af293 and CEA17Δ*akuB*^KU80^) underwent TEM analysis using a JEOL JEM-1400 electron microscope. TEM data for the CEA17Δ*akuB*^KU80^ sample was reproduced from previously published results (Chakraborty et al., 2021) for comparison with the other two strains. Fungal mycelia were treated with 2.5% glutaraldehyde, 2% paraformaldehyde, and 0.1 M cacodylate buffer, followed by embedding in 2% agarose to fix cellular organelles and prevent shrinkage. A secondary fixation with 0.1 M osmium tetroxide was performed. Dehydration was achieved using a series of acetone solutions with increasing concentrations, followed by infiltration with epoxy resins and acetone in proportions of 25:75, 50:50, and 75:25, respectively. Samples were incubated overnight in the 75:25 resin-acetone solution, then treated with 100% resin for two days with several resin changes. Finally, the samples were placed in an oven at 70°C to prepare the blocks. Ultrathin sections were cut using a LEICA EM UC7 microtome, stained with 1% uranyl acetate and lead acetate, and mounted on carbon-coated grids. TEM imaging focused on perpendicular cross-sections of the hyphae, with 100 measurements of cell wall thickness performed for each group (**Supplementary Table 2**). Cell wall thickness for all samples, including CEA17Δ*akuB*^KU80^, was measured using ImageJ software.

### ^13^C and ^15^N solid-state NMR

High-resolution solid-state NMR experiments were performed on either a Bruker Avance III 800 MHz (18.8 T) NMR spectrometer at the National High Magnetic Field Laboratory, Tallahassee, FL or a Bruker Avance Neo 800 MHz NMR at Michigan State University, East Lansing, MI. All ^13^C/^15^N detection experiments were performed using a 3.2 mm MAS probe with a MAS frequency of 10-15 kHz at ambient temperatures of 293-298 K. The ^13^C chemical shifts were calibrated externally to the tetramethylsilane (TMS) scale using the adamantane methylene carbon resonance at 38.48 ppm. The ^15^N chemical shifts were calibrated using the methionine amide resonance at 127.88 ppm, as observed in the model tripeptide N-formyl-Met-Leu-Phe-OH. Typical radiofrequency pulse field strengths used were 71.4-83.3 kHz for ^1^H hard pulses, decoupling, and during cross-polarization; 50-62.5 kHz for ^13^C pulses; and 41 kHz for ^15^N pulses. Experiments analyzing dynamics and hydration were carried out on a Bruker Avance III 400 MHz (9.4 T) spectrometer at Michigan State University, equipped with a 3.2 mm triple-resonance MAS probe, at temperatures ranging from 293-298 K, and 280 K for water-edited experiments. All experimental parameters are listed in **Supplementary Tables 3 and 4**.

### ^1^H solid-state NMR

^1^H-detection 2D hCH (Barbet-Massin et al., 2014) and 2D ^1^H-^13^C refocused INEPT-HSQC (Bodenhausen & Ruben, 1980) experiments were performed on all three *Aspergillus* samples using a Bruker Avance Neo 800 MHz NMR spectrometer in Michigan State University, equipped with a 1.6 mm triple-resonance Phoenix MAS probe. The samples were spun at 40 kHz MAS, with the temperature set to 290-300 K. DSS and D_2_O was added to measure the sample temperature and reference the ^1^H chemical shifts, with the DSS ^1^H peak set to 0 ppm. For 2D hCH, the radiofrequency pulse field strengths were 80 kHz for ^1^H during 90° pulses and cross-polarization (CP), and 40 kHz ^13^C during CP. The swept-low-power TPPM (slpTPPM) heteronuclear decoupling sequence was implemented with a field strength of 10 kHz on the ^1^H channel during the t_1_ evaluation period (Lewandowski et al., 2011). A WALTZ-16 heteronuclear decoupling sequence (Shaka et al., 1983) was applied with a field strength of 12 kHz on both ^13^C and ^15^N channels during the t_2_ acquisition period. To suppress water signals, the MISSISSIPPI (Zhou & Rienstra, 2008) pulse sequence was applied with a field strength of 20 kHz for 150-200 ms. 2D data were acquired using the States-TPPI quadrature detection method (Marion et al., 1989).

### 1D ssNMR experiments for screening carbohydrate dynamics

1D ^13^C spectra were acquired using four polarization methods to selectively detect signals from molecules with distinct dynamics. The *J*-coupling-based ^1^H-^13^C refocused Insensitive Nuclei Enhancement by Polarization Transfer (INEPT) experiment (Elena et al., 2005) targeted the most mobile molecules by using *J*-coupling to transfer magnetization between bonded ^1^H and ^13^C nuclei. This approach efficiently detects highly mobile molecules with long transverse relaxation times during four delays of 1/4*J*_CH_, 1/4*J*_CH_, 1/6*J*_CH_, and 1/6*J*_CH_ in the pulse sequence, where *J*_CH_ represents the carbon-hydrogen *J*-coupling constant and was set to 140 Hz. Second, 1D ^13^C direct polarization (DP) experiment with a short recycle delay of 2 s was employed to preferentially detect mobile molecules with fast ^13^C-T_1_ relaxation. Third, for quantitative detection, the same 1D ^13^C DP experiment was utilized, but with a very long recycle delay of 30 s and 35 s for *A. nidulans* (A28) and *A. fumigatus* (Af293), respectively. Lastly, the dipolar-mediated ^1^H-^13^C CP with 1 ms contact time was used to preferentially polarize rigid components. These diverse polarization methods facilitated the spectroscopic selection of different molecular dynamics within the samples, with zoomed spectra of the carbohydrate regions represented in **Supplementary Fig. 1**.

### 2D ssNMR for resonance assignment

Through-bond carbon connectivity was established using either scalar and dipolar-based polarization transfer techniques in the 2D ^13^C refocused *J*-INADEQUATE experiment (Cadars et al., 2007; Lesage et al., 1999). Similar to the polarization methods applied in the previous section for 1D experiments, two polarization schemes with 2 s time delays between scans were implemented for these 2D experiments: 2D ^13^C CP refocused *J*-INADEQUATE for detecting rigid molecules and ^13^C DP refocused *J*-INADEQUATE with 2 s recycle delays for detecting mobile components (**Supplementary Fig. 2**). Each of the four delays during the *J*-evolution period was set to 2.3 ms, optimized by tracking carbohydrate intensity. Through-space homonuclear ^13^C-^13^C correlations were recorded using the CP-based CORD sequence (Hou et al., 2013; Lu et al., 2015), where the mixing time was 50 ms for *A. nidulans* and *A. fumigatus* strains at 15 kHz MAS (**Supplementary Fig. 3**). Resonance assignments were further validated through cross-comparison with chemical shifts indexed in the Complex Carbohydrate Magnetic Resonance Database (Kang et al., 2020) as listed in **Supplementary Tables 5** and **6**.

### Molecular composition by intensity analysis

Molecular composition was evaluated by selecting analyzing the peak volumes of well-defined signals in 2D ^13^C spectra: CORD for the rigid portion and DP refocused *J*-INADEQUATE for the mobile fraction (**Supplementary Table 7**). In CORD spectra, quantification was achieved by calculating the mean of the resolved cross-peaks for each carbohydrate. In INADEQUATE spectra, exclusively well-differentiated spin connections were considered. The relative abundance of a specific polysaccharide was quantified by normalizing the sum of integrals with their respective counts, with standard errors calculated by dividing the standard deviation of the integrated peak volume by the total cross-peak counts; the overall standard error was then derived as the square root of the sum of the squared standard errors for each polysaccharide, as previously reported (Dickwella Widanage et al., 2024).

### MAS-DNP sample preparation and measurement of intermolecular interactions

The critical step in preparing fungal samples for MAS-DNP measurement involves incorporating stable biradicals and thoroughly mixing them with the sample. The biradical AMUPol (Sauvée et al., 2013) was mixed with a partially deuterated solvent of d_8_-glycerol/D_2_O/H_2_O (60/30/10 Vol%) to prepare a 10 mM stock solution. The inclusion of d_8_-glycerol acted as a cryoprotectant, and partial deuteration reduced proton density in the solvent, facilitating efficient ^1^H-^1^H spin diffusion from the solvent to the molecules of interest. The ^13^C,^15^N-labeled *A. nidulans* material was gently grounded using a mortar and pestle in the radical solution to ensure effective distribution of the radicals that can then diffuse into the fungal cell wall. This process ensures a uniform distribution of radicals within the sample, which leads to enhanced sensitivity in subsequent measurements. Approximately 30 mg of the ground sample was packed into a 3.2 mm sapphire rotor and subjected to MAS-DNP at a 10 kHz MAS frequency and 100 K.

To detect intermolecular interactions, we performed 2D ^15^N-^13^C and ^13^C-^13^C long-range correlation experiments using the processed *A. nidulans* sample on a 600 MHz/395 GHz MAS-DNP system at the National High Magnetic Field Laboratory (Dubroca et al., 2018). The typical radiofrequency field strengths for ^1^H, ^13^C, and ^15^N were 100 kHz, 50 kHz, and 50 kHz, respectively. The MAS frequency was set to 10 kHz. The DNP buildup time of the *A. nidulans* sample measured by saturation recovery was 2.8 s. Consequently, the recycle delays for all MAS-DNP experiments were set to 3.6 s (∼1.3 times the buildup time) for the highest signal-to-noise ratio within a given experimental time. The cathode current from the gyrotron was set at 150 mA and a voltage of 16.2 kV corresponding to ∼ 395.145 GHz and 12 W power at the probe base. The sensitivity enhancement factor (ε_on/off_) was measured by comparing the ^13^C signal intensity acquired with and without microwave (μw) irradiation (Chakraborty et al., 2020; Mentink-Vigier et al., 2015), and was found to be 27-fold.

For acquiring 2D ^15^N-^13^C heteronuclear correlation spectra, the NCACX pulse sequence (Baldus et al., 1998; Pauli et al., 2001) was employed. This sequence included a double-CP sequence with 0.5 ms of contact time for efficient polarization transfer from ^1^H-^15^N CP and 4 ms for ^15^N-^13^C CP. The ^15^N-^13^C CP was followed by a ^13^C-^13^C PDSD mixing period, with 0.1 s used for mapping short-range intramolecular cross peaks, and 3.0 s used for detecting both short-range intramolecular cross peaks and long-range intermolecular interactions, which occur on the sub-nanometer length scale. 2D ^13^C-^13^C homonuclear correlations were measured using the Proton-Assisted Recoupling (PAR) pulse sequence (De Paëpe et al., 2008; Donovan et al., 2017). A 2 ms PAR period was used for detecting short-range correlations, while a 20 ms PAR period was used for detecting long-range intermolecular cross peaks (**Supplementary Table 8**). The ^1^H and ^13^C irradiation frequencies for PAR were set at 56 kHz and 53 kHz, respectively. The number of scans was 8 for each 1D CP spectrum, 32 for 2D N(CA)CX, and 32 for 2D PAR.

### Solid-state NMR of polymer hydration and dynamics

All experiments investigating polymer hydration and dynamics were conducted on a Bruker Avance Neo 400 MHz (9.4 T) NMR spectrometer at Michigan State University using a 3.2 mm HCN MAS Bruker probe. The temperature was set to 280 K and 298 K for hydration and dynamics experiments, respectively. To assess the water accessibility of the polysaccharides, we employed 1D ^13^C and 2D water-edited ^13^C-^13^C correlation spectra (Ader et al., 2009; White et al., 2014). Initially, all protons were excited by applying a hard 90° pulse on the ^1^H channel, followed by a ^1^H-T_2_ filter to suppress the magnetization of proton resonances with ^1^H-T_2_ relaxation. Carbohydrates typically have substantially shorter ^1^H-T_2_ values than water, leading to the suppression of carbohydrate resonances and selectively retaining the proton magnetization originating from the mobile water. Subsequently, the proton polarization of water was transferred to nearby molecules, e.g., well-hydrated carbohydrates, through a ^1^H-^1^H mixing period. The polarization was then transferred to carbon nuclei through CP with a contact time of 1 ms for high-resolution ^13^C detection. Specifically, a ^1^H-T_2_ relaxation filter of 1.2 ms × 2 and 1.6 ms × 2 was used for Af293 and A28, respectively. This filter suppressed carbohydrate signals to less than 10% while retaining a minimum of 80% of water magnetization (**Supplementary Fig. 4**). For the 1D water-edited experiment, the ^1^H mixing time was systematically varied from 0 to 100 ms. These relative intensities were plotted as a function of the square root of the ^1^H mixing time, generating buildup curves for various carbon sites (**Fig. 5a**). For the 2D version of water-edited experiments, the ^1^H-^1^H mixing period was set to 4 ms and a 50 ms DARR mixing period was employed. The intensity ratios (S/S_0_) between both the water-edited spectrum (S) and a control 2D spectrum (S_0_) were analyzed, reflecting the water retention around each carbon site (**Supplementary Table 9**).

The dynamics of cell wall components were assessed via the analysis of ^13^C spin-lattice (T_1_) relaxation times. This was initially probed using a series of 2D ^13^C-^13^C correlation spectra with a variable z-filter period (0.1 s, 1 s, 3 s, and 9 s) (Wang et al., 2015), as illustrated in **Supplementary Fig. 5**. For ^1^H-T_1ρ_ relaxation measurement, the Lee-Goldburg (LG) spinlock sequence was utilized with varied ^1^H spinlock times ranging from 0.1 ms to 19 ms, resulting in 12 spectra (**Supplementary Fig. 6**). This experiment provided carbohydrate-specific information on polymer dynamics. The influence of ^1^H-^1^H dipolar couplings for ^1^H-T_1ρ_ relaxation measurements was suppressed by applying the LG block during the spinlock and CP period. The intensity of each peak was quantified, normalized by the number of scans, and fit using a single-exponential equation to obtain the relaxation time constants for different carbon sites (**Supplementary Figs. 5-6** and **Tables 10-11**).

## Results and Discussion

### Polysaccharides are dynamically distinct in A. nidulans *and* A. fumigatus cell walls

The dynamic profiles of fungal cell wall polysaccharides were rapidly screened through a series of 1D ^13^C experiments measured on *A. fumigatus* (Af293) and *A. nidulans* (A28). The most mobile molecules were identified using a *J*-coupling-based refocused INEPT experiment (**Fig. 1a**). The use of scalar coupling for polarization transfer from ^1^H to ^13^C and the lack of dipolar decoupling during the transfer period eliminated the signals of all rigid biomolecules characterized by strong ^1^H-^1^H dipolar couplings. Two types of ^13^C DP spectra measured with either a short recycle delay of 2 s for preferential detection of relatively mobile molecules with rapid ^13^C-T_1_ relaxation (**Fig. 1b**) or a long recycle delay of 35 s for quantitative detection of all molecules, ensuring unbiased observation by providing sufficient time for relaxation (**Fig. 1c**). Rigid molecules were identified using a dipolar-coupling-based ^13^C CP experiment (**Fig. 1d**). Five major structural polysaccharides and their relative mobility identified in *A. nidulans* are summarized in **Fig. 1e**.

**Figure 1.**
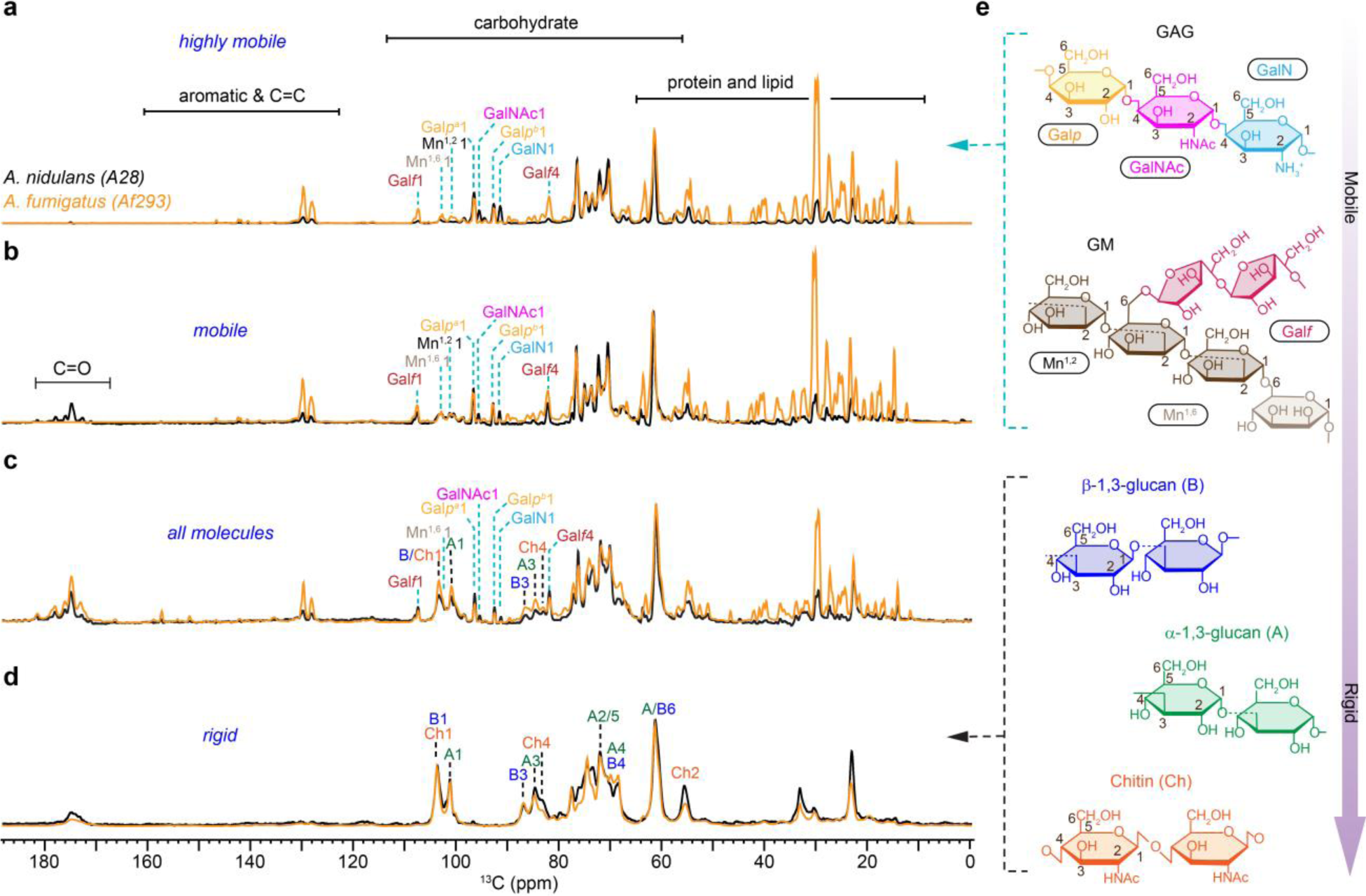
Dynamical gradient of polysaccharides in *Aspergillus* cell walls. From top to bottom are four sets of 1D ^13^C spectra measured with **a,** refocused INEPT experiment for probing the most dynamic molecules, **b**, DP spectra with short recycle delay of 2s for selection of mobile components. **c,** DP with long recycle delays for quantitative detection of all molecules. **d,** CP for selecting rigid polysaccharides. The spectra of *A. nidulans* (A28) and *A. fumigatus* (Af293) are shown in black and orange, respectively. For example, the Gal*f*1 peak at 107 ppm annotates the carbon 1 of glucofuranose (Gal*f*), which is the sidechain in the galactomannan (GM). Dash lines in cyan and black indicate the key peaks of mobile and rigid polysaccharides, respectively. Simplified structure representations are shown for key polysaccharides. **e,** Structural representation of key carbohydrate components following the dynamic gradient of an increasing level of rigidity from top to bottom as derived from the data only for *A. nidulans*. The NMR abbreviations for different polysaccharides and their monosaccharide units are labeled.

Galactosaminogalactan (GAG) is found to be highly dynamic in both *A. fumigatus* and *A. nidulans*, but its content, especially the amount of GalNAc and GalN residues has been reduced in *A. fumigatus*. The prominence of Gal*p*, GalNAc, and GalN signals in the INEPT spectra of both *Aspergillus* species (**Fig. 1a**) validated the highly dynamic nature of GAG, which comprises these three monosaccharide units. This carbohydrate polymer is typically found on the cell wall surface (Briard et al., 2020). This dynamic behavior is likely attributed to GAG’s limited interaction with the inner rigid core of the cell wall. Compared to *A. fumigatus*, *A. nidulans* showed significantly stronger C1 signals of GalNAc and GalN at 91 ppm and 95 ppm, respectively. These changes were consistently observed in the INEPT spectrum (**Fig. 1a**), as well as the two DP spectra measured with short and long recycle delays (**Fig. 1b, c**). These observations suggest an elevation in the surface charge of *A. nidulans* because GalN typically exists in its cationic unit (Fernando et al., 2023), GalNH_3_^+^, within fungal cells and therefore suggests variation in the physiochemical properties.

Galactomannan (GM) was found to be highly mobile in *A. fumigatus* but only partially dynamic in *A. nidulans*. This cell wall mannan consists of a linear backbone polysaccharide composed of a repeating tetramannoside oligosaccharide constituting of α-1,6 and α-1,2-linked mannose residues (Mn^1,2^ and Mn^1,6^) (Henry et al., 2016) along with a sidechain formed by galactofuranose residues (Gal*f*) (Henry et al., 2019). Tracking the signature signals of these sugar residues, for example, the Gal*f* carbon 1 (Gal*f*1) at 107 ppm and the mannose carbon 1 (Mn1) at 101-102 ppm, revealed that the GM content remained consistent in two *Aspergillus* species (**Fig. 1c**). Interestingly, GM exhibited high mobility in *A. fumigatus*, just like that of GAG, as indicated by its full intensity in the INEPT spectrum (**Fig. 1a**). In contrast, GM displayed only partial mobility in *A. nidulans*, with its signals predominantly appearing in the DP spectrum measured with 2-s recycle delays (**Fig. 1b**) but not in the INEPT spectrum (**Fig. 1a**). Despite its similar content in both *Aspergillus* species, GM has significantly reduced mobility in *A. nidulans*. Since GM is known to be covalently linked to β-1,3-glucan or β-1,3-glucan-chitin complex (Latgé, 2007), such interactions might have become more extensive in the cell walls of *A. nidulans,* which reduced the mobility of GM. Alternatively, the location of GM may differ between these two species, with this molecule potentially being less surface-exposed in *A. nidulans*. The zoomed spectra of the carbohydrate region ranging from 55-110 ppm is presented in **Supplementary Fig. 3**.

In both *A. fumigatus* and *A. nidulans*, the rigid components of their cell walls were primarily characterized by the prevalence of α-1,3- and β-1,3-glucans, alongside chitin (**Fig. 1d**). The only noticeable change, discernable within the limited resolution of a 1D spectrum, is the lower intensities of chitin peaks, such as the carbon 2 (Ch2) at 55 ppm and carbon 4 (Ch4) at 83 ppm. Therefore, *A. fumigatus* has a lower content of chitin in its cell wall.

Peaks corresponding to α-1,3- and β-1,3-glucans, primarily situated within the rigid domains, were also observed in the INEPT and 2s DP spectra designed for detecting the mobile molecules, albeit with low intensities (**Fig. 1a, b**). These weak peaks include the A1 at 101 ppm, A3 at 84 ppm, and B3 at 87 ppm. The observed dynamic variation is a consequence of the widespread distribution of these glucans across the cell wall, where they serve versatile functions in reinforcing both the rigid structural components and the flexible matrix of the cell wall. Compared to *A. nidulans*, *A. fumigatus* also showed 4-times stronger signals of protein and lipid, two components primarily residing in the mobile fractions (**Fig. 1a-c**). These protein and lipid signals may have various sources, as discussed recently (Gautam et al., 2024), and are therefore not the focus of this study, but their roles could be worth investigating in the future through extraction or removal procedures such as SDS treatment (Ehren et al., 2020).

### Molecular composition of the mobile and rigid portion

Well-resolved carbohydrate signals were identified using high-resolution 2D ^13^C-^13^C correlation spectra. Resonance assignment was achieved mainly using the through-bond refocused ^13^C INADEQUATE experiment, which allows us to unambiguously track the carbon connectivity within each carbohydrate unit, thus resolving the ^13^C chemical shifts of each carbon site (**Fig. 2a, b**). The resulting spectrum is asymmetric, correlating single-quantum (SQ) chemical shift with double-quantum (DQ) chemical shift, which is the sum of two SQ chemical shifts from two directly bonded carbons. These experiments were conducted separately for the rigid and mobile fractions using CP and DP for initial polarization, respectively.

**Figure 2.**
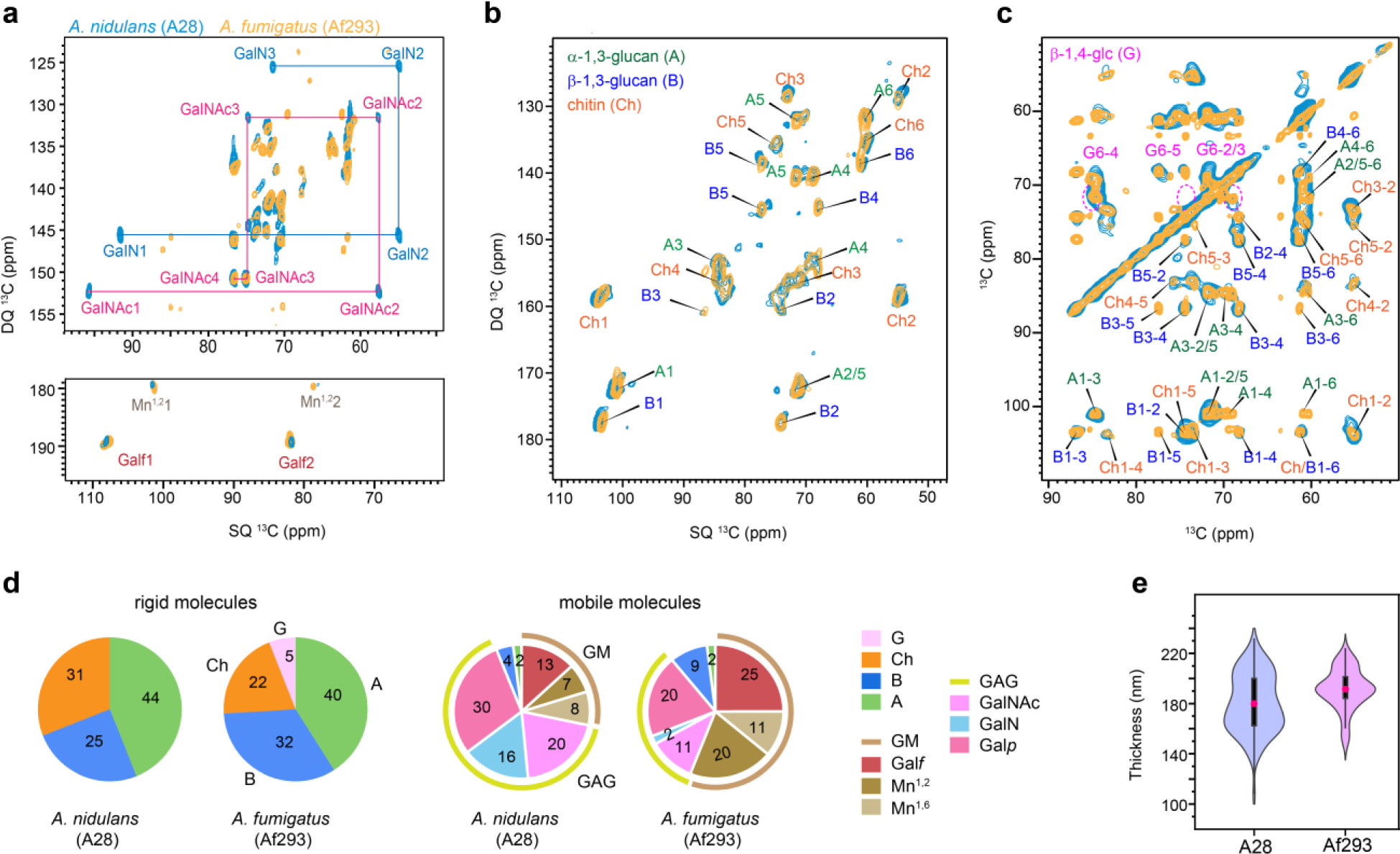
Molar composition of polysaccharides in rigid and mobile domains. **(a)** Mobile components detected by 2D ^13^C DP refocused *J*-INADEQUATE spectra of *A. nidulans* A28 strain (turquoise) and *A. fumigatus* A293 strain (yellow). The full names and NMR abbreviations are listed for key monosaccharide units and polysaccharides. **(b)** Rigid components detected by 2D ^13^C CP refocused *J*-INADEQUATE spectra of *A. nidulans* (turquoise) and *A. fumigatus* (yellow). **(c)** 2D ^13^C-^13^C CORD mixing correlation spectra measured showing signals of rigid polysaccharides in *A. nidulans* (turquoise) and *A. fumigatus* (yellow). Dash line circles in magenta highlight the signals of β-1,4-glucose units, which are observed in *A. fumigatus* but missing in *A. nidulans*. **(d)** Molar compositions of polysaccharides in the rigid (left) and mobile (right) fractions of the two *Aspergillus* strains. The values were calculated using the peak volumes in 2D CORD and DP *J*-INADEQUATE spectra. NMR abbreviations are given for key residues: B: β-1,3-glucan; Ch: chitin; A: α-1,3-glucan; G: β-1,4-linked glucopyranose residues; GM: galactomannan; GAG: galactosaminogalactan; Mn^1,2^: 1,2-linked mannose; Mn^1,6^: 1,6-linked mannose. Gal*f*, GalN, Gal*p* and GalNAc are standard abbreviations of sugar residues. **(e)** Violin plots depict the distribution of 100 measurements based on TEM images, with a minimum of 10 cells analyzed for each sample of *A. nidulans* (A28) and *A. fumigatus* (Af293). In each violine plots, the black rectangle represents the interquartile range (25-75% IQR) in ascending order and the pink circle denoted the mean of the dataset while the black vertical line denotes the standard range of the 1.5IQR.

In the mobile fraction of both *A. nidulans* A28 and *A. fumigatus* Af293, we successfully resolved all carbon sites for the monosaccharide units present in GAG and GM (**Fig. 2a**), and the glucose units forming α- and β-1,3-glucans (**Supplementary Fig. 2**). For instance, the GalNAc and GalN residues in GAGs were tracked through the distinctive signals of C2 with unique SQ chemical shifts of 55 and 57 ppm. These chemical shifts are specific to the carbon site covalently linked to nitrogen. GalNAc1 and GalN1 further correlate with the carbon 1 sites (SQ chemical shift of 92 and 96 ppm), resonating at DQ chemical shifts of 147 and 153 ppm. Similarly, in the case of Gal*f*, the sidechain of GM, its carbon 1 (Gal*f*1) and carbon 2 (Gal*f*2) resonate at SQ chemical shifts of 107 and 82 ppm, respectively, resulting in a DQ chemical shift of 189 ppm (**Fig. 2a**). The observations of GAG and GM, along with a minor presence of α- and β-1,3-glucans, support their prominent roles in the mobile domains of cell walls, which encompass the outer surface and the soft matrix, in both *Aspergillus* species.

Both *A. nidulans* and *A. fumigatus* exhibited consistent presence of chitin, α-1,3-glucan and β-1,3-glucan in the rigid fraction (**Fig. 2b**). Strong signals for β-1,4-glucose residues were exclusively detected in the *A. fumigatus* Af293 cell wall as shown by the 2D ^13^C CORD spectrum, while they were not observed in *A. nidulans* (**Fig. 2c** and **Supplementary Fig. 3**). These β-1,4-glucose residues are part of the mix-linked β-1,3/1,4-glucan typically found as a linear terminal domain in *A. fumigatus*. The absence of their signals in *A. nidulans* indicates a lack of such structural domains within its β-glucan matrix.

Upon checking the gene sequence for the absence of β-1,3/1,4-glucan synthase, encoded by the gene *tft1* (Afu3g03620) and the protein XP748682 in *A. fumigatus,* a BLAST search on NCBI and VEuPathDB confirmed that this protein is present in many *Aspergillus* species, as corroborated by (Samar et al., 2015), but absent in the *A. nidulans* strain FGSC 8444, which is the only strain of *A. nidulans* sequenced so far. The absence of β-1,3/1,4-glucan has not been shown earlier. A cellulase gene, *celA* (AN8444), with putative functions involved in β-1,3/1,4-glucan synthesis, has been recently analyzed in *A. nidulans*. However, the absence of β-1,4/1,3 glucan has not been checked in the *celA* mutant and the Blast for the entire *celA* gene and the *A. fumigatus* Tft1 protein.

These spectroscopic observations were numerically presented as the molar composition (**Fig. 2d** and **Supplementary Table 7**), determined by averaging the peak volume of resolved signals within each monosaccharide type and the polysaccharide formed by these units. Both A28 and Af293 contained approximately 40% of α-glucans in their rigid fractions, but the chitin content was higher in *A. nidulan*s, accounting for 31% of its rigid polysaccharides, compared to Af293 (22%) and CEA17Δ*akuB*^KU80^ (8%) (Chakraborty et al., 2021). *A. nidulans* mainly contains β-1,3-glucan, making up 25% of the rigid fraction, while Af293 contains 32% of β-1,3-linked glucose units along with 5% of β-1,4-linked glucose residues. Assuming a 1:1 molar ratio of its mixed linkages, the content of β-1,3/1,4-glucan in *A. fumigatus* Af293 is estimated to be 10%, leaving 28% of the rigid portion as β-1,3-glucan. Hence, there are major structural differences in the chitin and β-glucan matrix between *A. nidulans* and *A. fumigatus* observed in the rigid core of the cell wall, although they exhibited highly comparable cell wall thickness (**Fig. 2e**).

In *A. fumigatus* Af293, the mobile fraction primarily consists of GM (56%) with a smaller amount of GAG (one-third), whereas in *A. nidulans*, GM content is lower (approximately 30%) and GAG content is higher, accounting for about two-thirds of the soft molecules (**Fig. 2d**). This contradicts previous biochemical results, which showed that *A. fumigatus* typically secretes more GAG with a higher content of GalNAc/GalN compared to Gal*p* than *A. nidulans* (A26), correlating with the higher virulence of *A. fumigatus* (M. J. Lee et al., 2015). This discrepancy may be attributed to differences in strains, as well as the distinct media, culture conditions, and durations used in these studies.

The structural feature of chitin, which was previously observed in *A. fumigatus* as a highly polymorphic carbohydrate polymer (Fernando et al., 2021), is also valid in *A. nidulans*. DNP-enhanced 2D ^13^C-^13^C and ^15^N-^13^C correlation spectra provided a clear view of all carbon sites and amide nitrogen in chitin (**Fig. 3a**), with all ^13^C and ^15^N chemical shifts documented in **Supplementary Tables 5 and 6**. Notable peak multiplicity was observed for most chitin signals, even at the cryogenic temperatures used in DNP. Significant examples include cross peaks involving C1, C4, and C6, where variations in C1 and C4 chemical shifts reflect torsional flexibility around the glycosidic linkage, and C6 variations demonstrate the hydroxymethyl’s structural flexibility. Such structural variations are also seen in cellulose and xylan in plant cell walls (Kirui et al., 2022; Phyo et al., 2018; Simmons et al., 2016). Room-temperature spectra offered better resolution, allowing differentiation of four types of chitin molecules (**Fig. 3b**). Additionally, multiplicity was observed for α-glucan, with C1 chemical shifts at 101 and 100 ppm for the major and minor forms, respectively (**Fig. 3b**). Type-a α-1,3-glucan is typically found in large quantities in *A. fumigatus*, while type-b usually contributes only about 2%, but increases to 10-20% of the entire cell wall when *A. fumigatus* is exposed to echinocandins (Dickwella Widanage et al., 2024).

**Figure 3.**
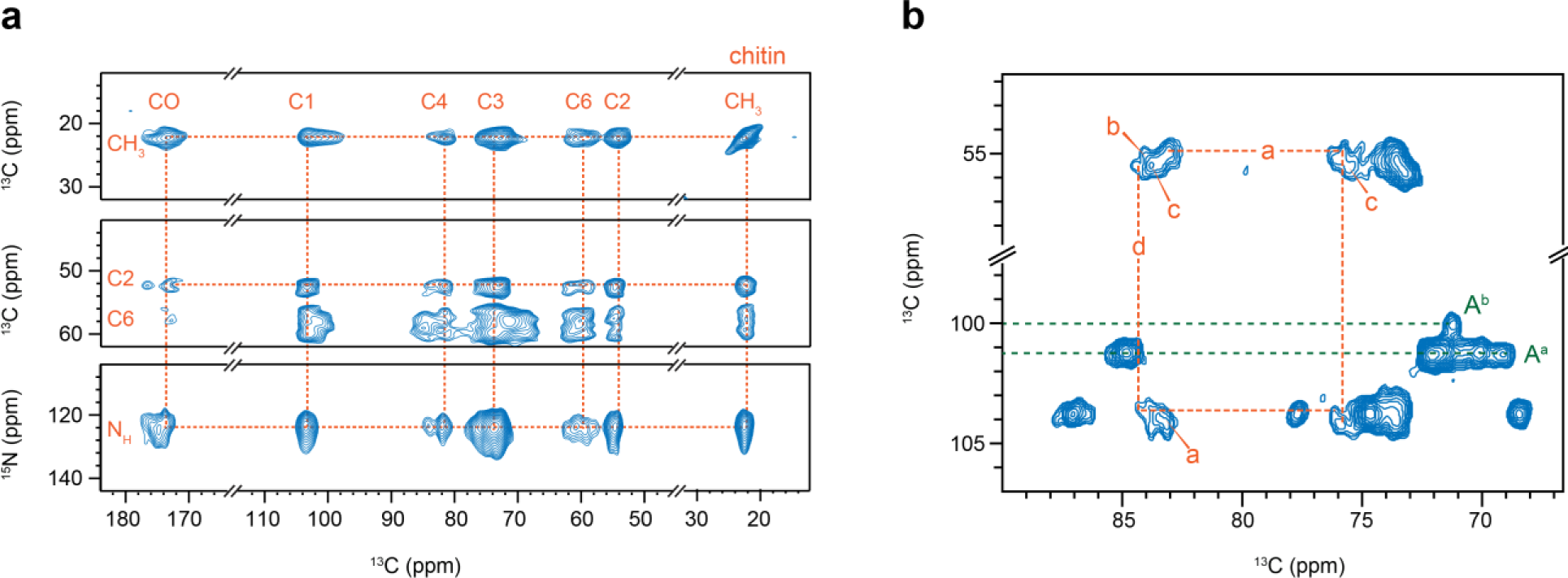
Structural complexity of chitin and α-glucan in *A. nidulans*. **(a)** Chitin signals resolved by 2D ^13^C-^13^C PAR correlation spectrum (top and middle) and 2D ^15^N-^13^C correlation spectrum (bottom panel). These spectra were measured using DNP for sensitivity enhancement. **(b)** Peak multiplicity observed in 2D ^13^C-^13^C CORD correlation spectrum. Dashed lines in orange show the carbon connectivity of type-a and type-d chitin. Dashed lines in green show the signals of type-a (A^a^) and type-b (A^b^) α-1,3-glucan.

### Proton-detected spectra of Aspergillus species

Proton detection offers a more time-efficient alternative for studying biomolecules (Marchand et al., 2022), and has recently been applied to the analysis of cell walls in fungi, plants, and bacteria (Bahri et al., 2023; Bougault et al., 2020; Duan & Hong, 2024; Ehren et al., 2020; Phyo & Hong, 2019; Safeer et al., 2023; Vallet et al., 2024; Yuan et al., 2021). We detected the rigid and mobile molecules within the *Aspergillus* cell walls using polarization transfer methods based on dipolar and scalar couplings, respectively (**Fig. 4a, b**). 2D hCH spectra exhibited analogous patterns in *A. nidulans* and both strains of *A. fumigatus* (**Fig. 4a**), reflecting the similarity in polysaccharide structure within their rigid fractions observed in ^13^C-based data (**Fig. 2b**), except for compositional changes. The J-coupling-based INEPT-HSQC spectra detected mobile molecules, where *A. nidulans* demonstrated significantly fewer peaks in the lipid/protein region (10-55 ppm) than *A. fumigatus* strains (**Fig. 4b**), consistent with the alteration of the molar fraction within the mobile fraction as shown in **Fig. 2a, d**.). INEPT-HSQC spectrum also exhibited well resolved signals of the terminal residues, such as the non-reducing ends of β- and α-glucans (**Fig. 4c, d** and **Supplementary Table 12**). The signals of these terminal residues were consistently observed in both *A. nidulans* and the Af293 strain of *A. fumigatus*, indicating similar chain lengths of their glucans. However, their signals became significantly weaker in *A. fumigatus* CEA17Δ*akuB*^KU80^, indicating different structure, presumably longer β-glucans in this model strain.

**Figure 4.**
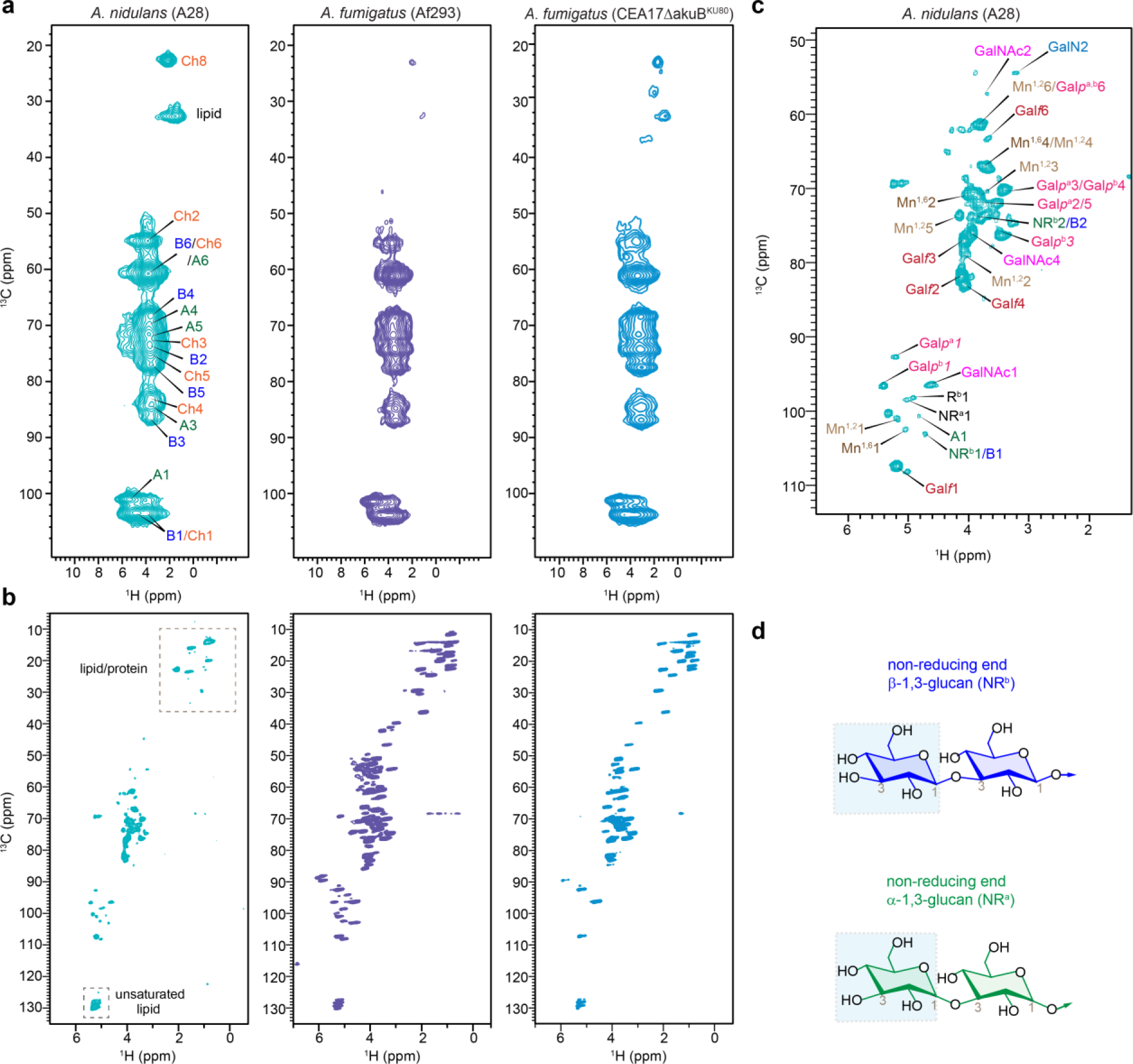
Carbohydrate structure from proton-detected solid-state NMR. **(a)** Rigid molecules detected in 2D ^13^C-^1^H hCH spectra of *A. nidulans* (A28), *A. fumigatus* (Af293), and *A. fumigatus* (CEA17Δ*akuB*^KU80^). (**b**) Mobile polysaccharides detected using 2D ^13^C-^1^H refocused *J*-INEPT-HSQC correlation experiment. (**c**) Zoomed-in view of carbohydrate region of *J*-INEPT-HSQC spectrum with resonance assignment. (**d**) Representative structure of non-reducing ends of α- and β-glucans are shown. All spectra were measured on an 800 MHz spectrometer at 40 kHz MAS. The experimental time is 3.4 h for each 2D hCH spectrum and 10.2 h for each J-INEPT-HSQC spectrum.

**Figure 5.**
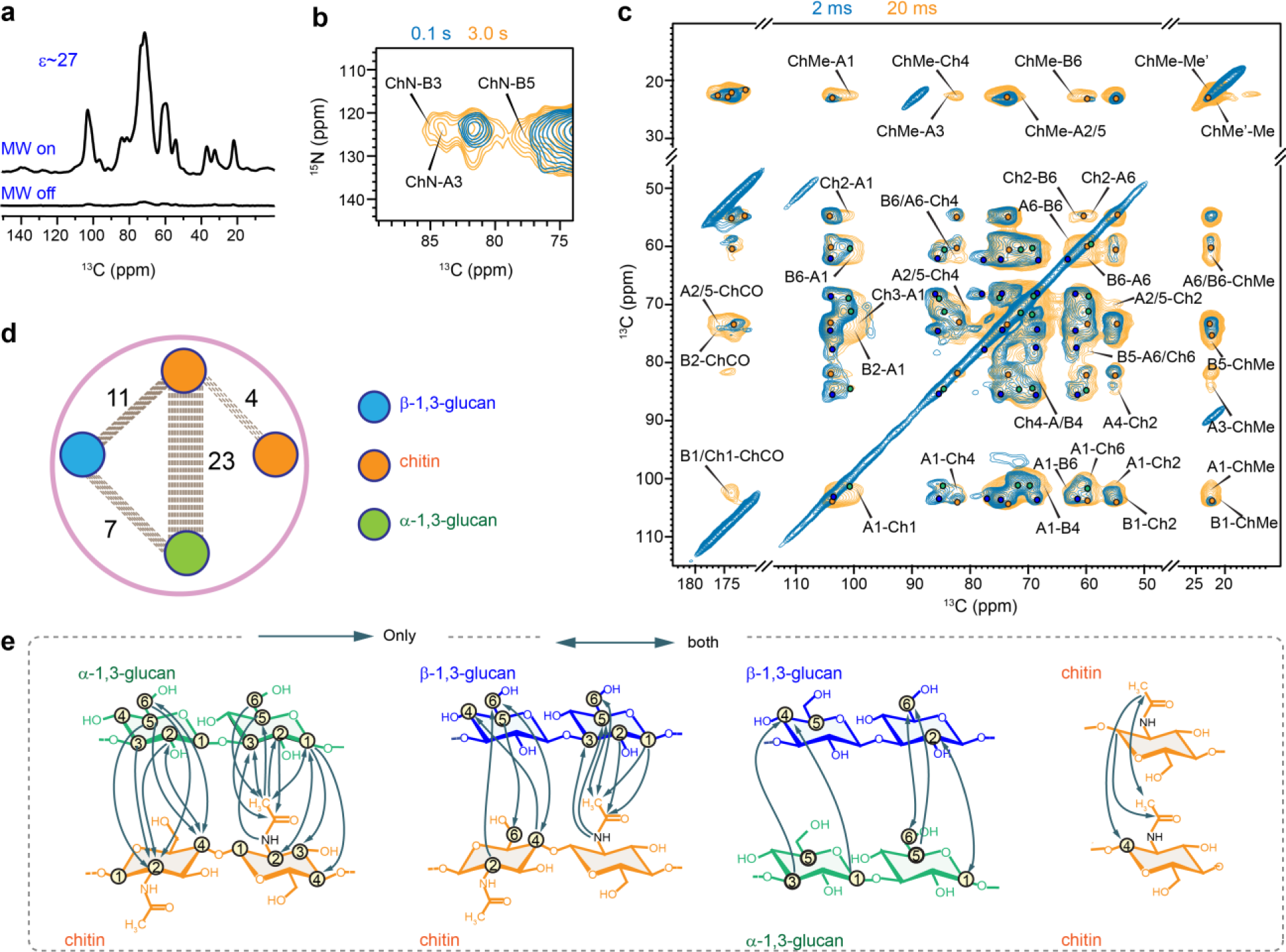
DNP-supported view of intermolecular interactions of *A. nidulans* polysaccharides. **(a)** An DNP enhancement of 27-fold was achieved on *A. nidulans* when comparing the spectra measured with and without microwave (MW). (b) Overlay of 2D ^15^N-^13^C correlation spectra measured with short (0.1 s; turquoise) and long (3.0 s; yellow) ^13^C-^13^C mixing periods **(c)** Overlay of two 2D ^13^C-^13^C correlation spectra measured with 20 ms (yellow) and 2 ms (turquoise) PAR mixing periods. Labels are provided only for the long-range intermolecular cross peaks uniquely present in the 20 ms PAR spectrum. (**d**) Overview of intermolecular cross peaks detected among different polysaccharides: β-glucans (blue), α-glucans (green) and chitin (orange). The dash lines represent the number of intermolecular interactions between the glucans. **(e)** Structural summary of intermolecular interactions observed in *A. nidulans*. The NMR polarization-based interactions have directionality as shown using arrow heads. For example, a cross peak may be observed from the C3 of α-1,3-glucan to the methyl of chitin (A3-ChMe), or vice versa (ChMe-A3).

It should be noted that proton detection has substantially shortened the experimental time to 3-10 hours per spectrum, compared to the 20-30 hours per spectrum for ^13^C detection, thus providing a rapid option for screening molecular composition of the fungal samples. The ^1^H linewidth is 0.09-0.13 ppm (70-110 Hz) and 0.66-1.3 ppm (530-1070 Hz) for mobile and rigid molecules even under the moderately low MAS of 40 kHz on moderately high field of 800 MHz in the experimental condition used here. For *A. nidulans*, the ^1^H linewidths were measured as 72 Hz for Gal*p*^b^1, 83 Hz for both Mn^1,2^5 and Gal*f*1 in the INEPT-HSQC spectrum, and 529 Hz for Ch8, 938 Hz for B4, and 1070 Hz for both Ch2 and Ch4 in hCH spectrum. The resolution will be substantially improved, especially for the rigid components, at faster MAS and higher magnetic fields (Safeer et al., 2023; Vallet et al., 2024).

### Packing of cell wall polysaccharides in A. nidulans

To study the intermolecular arrangement of polysaccharides in *A. nidulans* cell walls, we used 2D ^13^C-^13^C and ^13^C-^15^N correlation experiments with extended mixing times, enabling polarization transfer at the sub-nanometer scale. MAS-DNP enhanced NMR sensitivity by 27-fold (**Fig. 5a**), facilitating the detection of weak intermolecular cross peaks between polysaccharides. 2D ^15^N-^13^C correlation spectra (**Fig. 5b**), obtained via the N(CA)CX experiment with varying mixing times, revealed strong cross peaks after long mixing (3s) between chitin amide nitrogen and carbons 3 and 5 of β-1,3-glucan (ChN-B3 and ChN-B5), as well as carbon 3 of α-1,3-glucan (ChN-A3).

The 2D ^13^C-^13^C spectrum, acquired with a 20 ms Proton Assisted Recoupling (PAR) period (De Paëpe et al., 2008; Donovan et al., 2017), displayed numerous long-range cross peaks indicating four types of intermolecular interactions (**Fig. 5c**). Firstly, significant cross peaks were observed between chitin and α-1,3-glucan, specifically between the chitin methyl carbon and the carbon 1, 3, and 2/5 sites of α-1,3-glucan (ChMe-A1, ChMe-A3, ChMe-A2/5), and between the chitin carbon 2 and the carbon 1 and 6 of α-1,3-glucan (Ch2-A1, Ch2-A6). Secondly, chitin showed cross peaks with β-1,3-glucan, such as ChMe-B6 and Ch4-B4. Thirdly, fewer cross peaks were detected between β- and α-1,3-glucans, including B6-A1 and A6-B6. Lastly, cross peaks between chitin methyl groups in magnetically inequivalent forms (ChMe-Me’ and ChMe’-Me) were identified. Detailed summaries of the identified intermolecular interactions are provided in **Fig. 5d, e**, and **Supplementary Table 8**.

It is noteworthy that 23 out of 45 observed intermolecular cross peaks were between chitin and α-1,3-glucan (**Fig. 5d**), supporting the concept that α-glucans extensively interact with chitin microfibrils in the rigid core of the mycelial cell wall. This concept, initially identified in *A. fumigatus* (Kang et al., 2018), is now confirmed in *A. nidulans*. Meanwhile, β-glucans exhibit moderate interactions with both chitin and α-glucan, indicating their loose packing within the structural core. This arrangement is likely reinforced by covalent linkages with chitin, as previously determined by chemical analyses (Latgé, 1999, 2007).

### Dynamics and water association

The water retention properties of cell wall polymers were analyzed using a water-editing experiment, as shown in **Supplementary Fig. 4** (Ader et al., 2009; White et al., 2014). In this method, a water-edited 2D ^13^C-^13^C correlation experiment was performed, utilizing a ^1^H-T_2_ relaxation filter to remove polysaccharide signals, followed by the transfer of water magnetization to the polysaccharides to detect the carbons proximal to water (**Fig. 6a**). The intensity ratio (S/S_0_) was determined for each resolvable carbon site, comparing the water-edited (S) and control (S_0_) spectra, as summarized in **Fig. 6b** and **Supplementary Table 9**. Both fungal species displayed relatively high S/S_0_ ratios (above 0.3) for β-1,3-glucans, indicating their significant role in maintaining the soft matrix and regulating water accessibility. In contrast, chitin and α-1,3-glucan exhibited lower S/S_0_ values (0.20-0.25), suggesting reduced water accessibility due to their physical association and the formation of larger, less permeable polymer domains (**Fig. 6b**). This hydration heterogeneity in *A. nidulans* cell walls is consistent with findings in *A. fumigatus* Af293, as shown here, and CEA17Δ*akuB*^KU80^ and RL578, as previously reported (Chakraborty et al., 2021; Kang et al., 2018). This structural feature is a conserved characteristic within *Aspergillus* cell walls.

**Figure 6.**
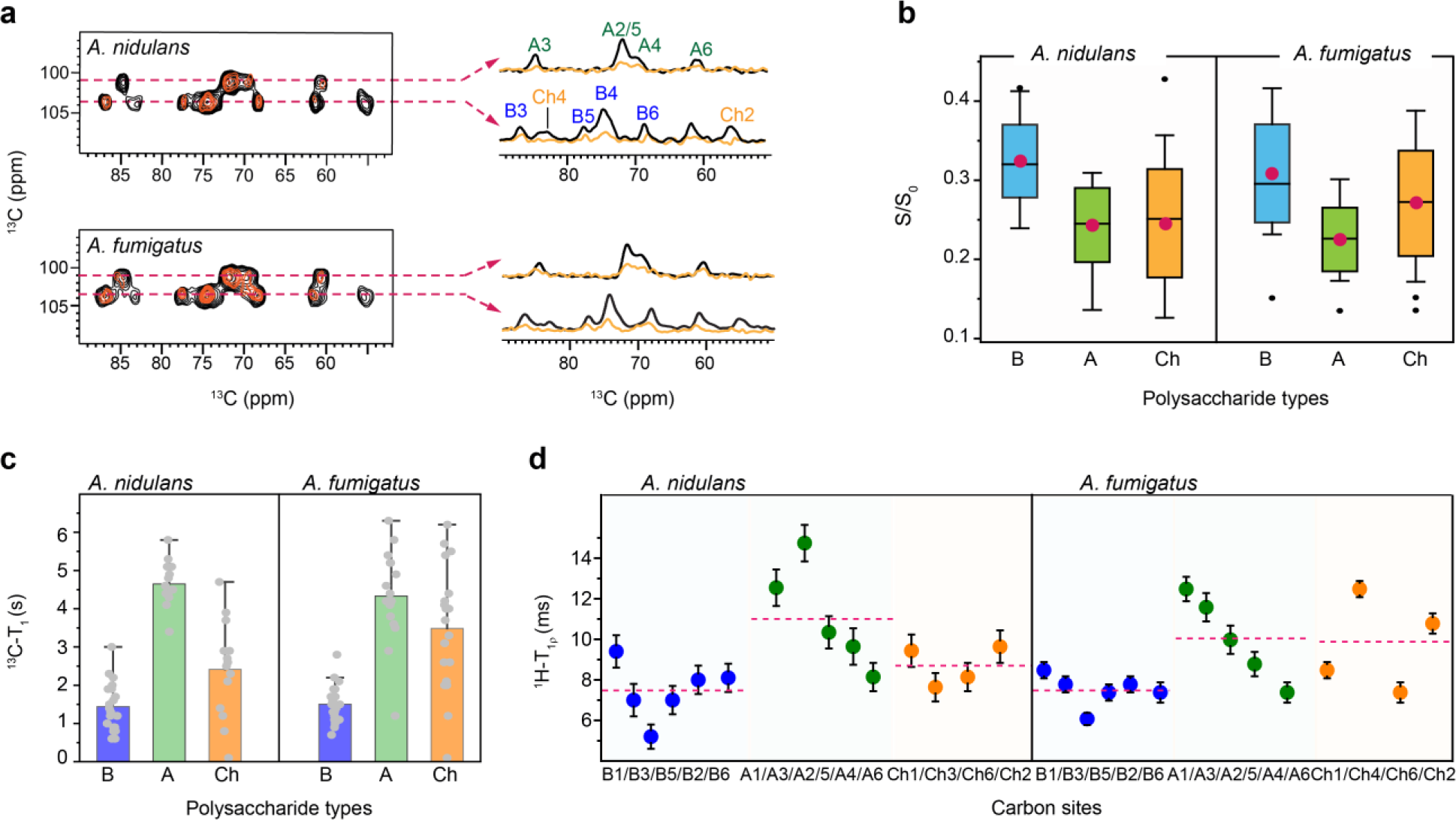
Hydration and the dynamics of *Aspergillus* polysaccharides. Data of water association and dynamics were compared between *A. nidulans* A28 strain and *A. fumigatus* Af293 strain. **(a)** Overlay of water-edited 2D ^13^C-^13^C spectra (orange) and control spectra (black). Cross sections were extracted at 101 ppm for α-1,3-glucan, C1 at 104 ppm for chitin, and β-1,3-glucan C1. **(b)** The average representation of intensity ratio of water edited spectra for *A. nidulans* and *A. fumigatus* where the glucans are color-coded encoded in a box the red circle represents the mean and the middle line represents the median while the bar with the cap represents the range and the black circle represents the outliers. **(c)** 2D ^13^C-T_1_ relaxation time constants measured for specific polysaccharide types encoded in Box representing the mean ± s.d. and whisker plotting with blue, green, and orange color for β-1,3-glucan (n=19 and 19), α-1,3-glucan (n= 14 and 15) and chitin (14 and 16). The average value of each polysaccharide type is represented in an open circle and the dark represents the outlier. **(d)** Site-specific ^1^H-T_1ρ_ relaxation time constants plotted against different carbon sites in β-1,3-glucan (B; blue, n=6), α-1,3-glucan (A; green, n= 5), and chitin (Ch; orange, n= 5). The average is shown in the solid and the dash lines.

To map out polysaccharide dynamics, relaxation experiments were conducted using ^13^C-T_1_ and ^1^H-T_1ρ_ techniques (**Supplementary Figs. 5, 6** and **Tables 10-11**). 2D ^13^C-T_1_ relaxation experiments were utilized to examine the dynamic behavior of polysaccharides on the nanosecond (ns) timescale. α-glucans had the slowest ^13^C-T_1_ relaxation, with 4.7 s in A28 and 4.3 s in Af293. For β-1,3-glucan, the ^13^C-T_1_ time constants were 1.4 s and 1.5 s, while for chitin, they were 2.4 s and 3.5 s for *A. nidulans* and *A. fumigatus* Af293, respectively (**Fig. 5c**). Similar trends were observed for ^1^H-T_1ρ_ data that probes motions happening on the microsecond timescale. For *A. nidulans*, the average relaxation times for β-1,3-glucan, α-1,3-glucan, and chitin were 7.5 ms, 11.0 ms, and 8.7 ms, respectively, while in the Af293 strain, the respective relaxation times were 7.5 ms, 10.1 ms, and 9.8 ms. The consistently long relaxation times for α-1,3-glucan across both nano-and millisecond timescales in *A. nidulans* and *A. fumigatus* (**Fig. 5c, d**) indicate that this carbohydrate polymer is spatially restricted, likely due to its dense packing with chitin microfibrils. This packing further limits water association, as shown in **Fig. 5b**. In contrast, the rapid relaxation of β-1,3-glucan reflects its high level of water association, underscoring its important role in maintaining the cell wall matrix.

### Conclusions and Perspectives

High-resolution solid-state NMR data of *A. nidulans* and *A. fumigatus* revealed only subtle compositional differences. *A. nidulans* showed lower levels of GAG, protein, and lipid in the mobile fraction and higher chitin content in the rigid fraction. Additionally, GM in *A. nidulans* is only partially dynamic, unlike the fully dynamic nature of GM in *A. fumigatus*. The glucan matrix in *A. nidulans* has also been restructured, with a predominance of β-1,3-glucans lacking terminal β-1,3/1,4-glucan domains. This also confirmed the lack of structural role of the β-1,3/1,4-glucan as shown in *A. fumigatus* with the *tft1* mutant (Samar et al., 2015). The similar polysaccharide composition suggests that the pathophysiological differences between the two species cannot be directly attributed to their cell wall composition (Gresnigt et al., 2018; Sugui et al., 2014). This also underscores the need for further research into protein and lipid components, which have been shown to be embedded in the polysaccharide matrix (Kniemeyer, 2011).

Despite the compositional differences, both *A. nidulans* and *A. fumigatus* exhibit highly comparable cell wall architecture, including thickness, dynamics, water association, and polysaccharide packing. This suggests that both *Aspergillus* species employ similar physical principles in their cell wall construction and confirms that the cell wall polymers serve the same biological functions in both species. Recent studies of *A. fumigatus* mycelial cell walls have identified rigid scaffolds formed by chitin, β-glucans, and an unexpected presence of α-1,3-glucan (Kang et al., 2018). Both α- and β-glucans are found in rigid and mobile phases, supporting the rigid core and forming the soft matrix (Chakraborty et al., 2021), while GM and GAG are primarily located in the mobile fraction, with GM chemically linked to β-glucan and β-glucan-chitin complexes (Latgé, 2007). This study has extended these biophysical insights to another important *Aspergillus* species.

It is important to recognize that the fungal cell wall is a highly dynamic structure, continuously reshuffling its composition and nanoscale organization in response to the fungus’s age, growth conditions, and environmental stressors (Gow & Lenardon, 2023). For *A. fumigatus*, recent solid-state NMR results have revealed that echinocandin treatment induces hyperbranched β-glucan formation, increases chitin and chitosan content, and creates new forms of semi-dynamic α-1,3-glucan, leading to a stiffer, less permeable, and thicker cell wall (Dickwella Widanage et al., 2024).

Similar remodeling has been observed in another *Aspergillus sydowii* under hypersaline conditions, suggesting that such cell wall reinforcement strategies are widespread among fungi to enhance survival under adverse conditions (Fernando et al., 2023). In addition, such structural schemes and remodeling mechanisms observed in mycelia do not apply to conidia: in dormant conidia, α-1,3-glucan and β-1,3-glucan are confined to the inner wall and shielded by RodA rodlets, with swelling disrupting this layer to enhance water access, while during germination, galactosaminogalactan appears in the mobile phase and chitin is incorporated into the inner wall (Lamon et al., 2023). While the current data identified a highly similar cell wall architecture in *A. fumigatus* and *A. nidulans,* it remains uncertain whether these structures respond differently to stress. Future solid-state NMR studies should also focus more on conidia, which is the infective propagule, as previous research has highlighted the presence of significant difference between the mycelia and conidia of *Aspergillus* (Latgé et al., 2017).

## CRediT authorship contribution statement

**Isha Gautam:** Writing – original draft, Investigation, Formal analysis. **Jayasubba Reddy Yarava:** Writing – review & editing, Investigation, Formal analysis. **Yifan Xu:** Writing – review & editing, Investigation. **Reina Li:** Writing – review & editing, Investigation. **Frederic Mentink-Vigier:** Investigation. **Faith J. Scott:** Investigation. **Michelle Momany**: Writing – review & editing, Resources. **Jean-Paul Latgé:** Writing – review & editing, Conceptualization. **Tuo Wang**: Writing – review & editing, Conceptualization, Funding acquisition.

## Acknowledgment

This work was supported by the National Institute of Health (NIH) grant AI173270. A portion of this work was performed at the National High Magnetic Field Laboratory, which is supported by National Science Foundation Cooperative Agreement No. DMR-2128556 and the State of Florida. The MAS-DNP system at NHMFL is funded in part by NIH and NIH RM1-GM148766. FJS acknowledges support by a Postdoctoral Scholar Award from the Provost’s Office at Florida State University. The authors thank Arnab Chakraborty and Liyanage Fernando for initial data analysis.

