## Supplementary Material for "Comparative Analysis of Polysaccharide and Cell Wall Structure in *Aspergillus nidulans* and *Aspergillus fumigatus* by Solid-State NMR"

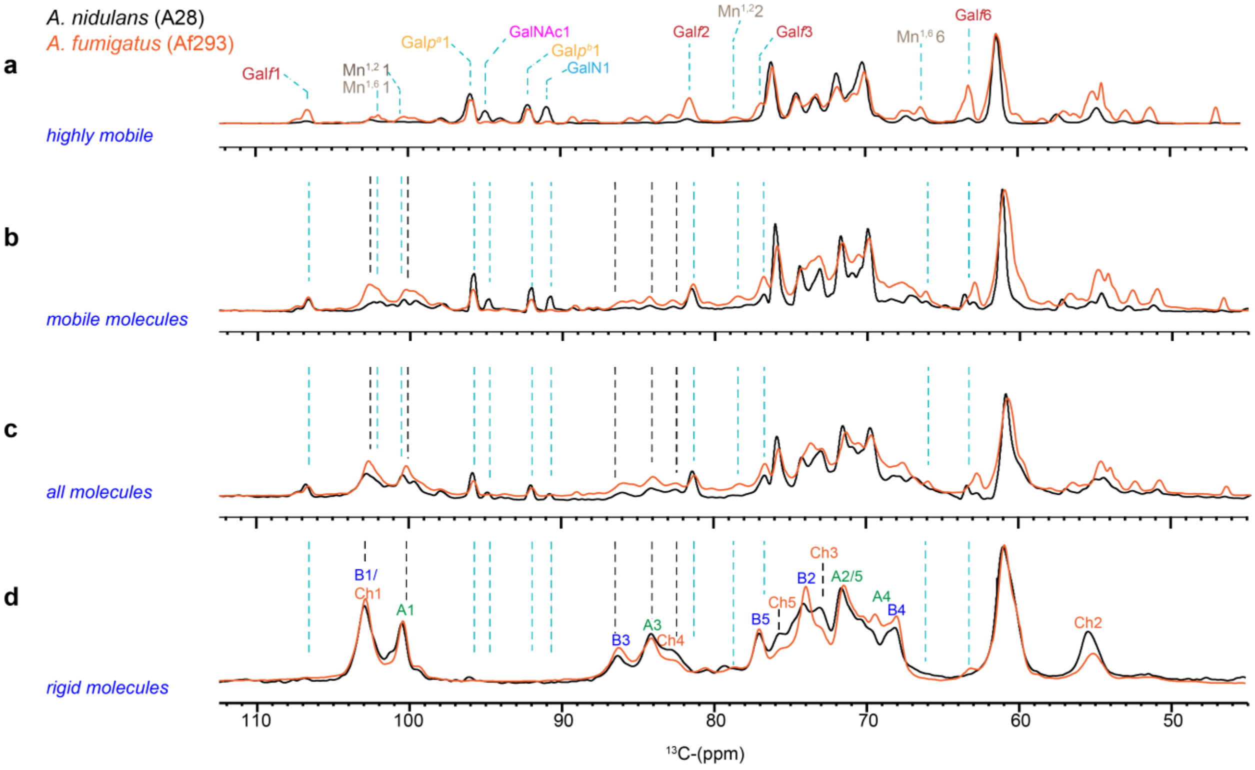


**Supplementary Figure 1.** Overlay for showing the zoom spectra of carbohydrate region. The spectra are color coded to show the black spectra for *A. nidulans* and orange for *A. fumigatus*. (**a**) Carbohydrates showing the highly mobile glucans obtained by INEPT experiment. (**b**) The semi mobile glucans probed by 2 s DP spectra and (**c**) The quantitative glucans represented by the 35 s DP spectra (**d**) The rigid molecules probed by CP experimentation.

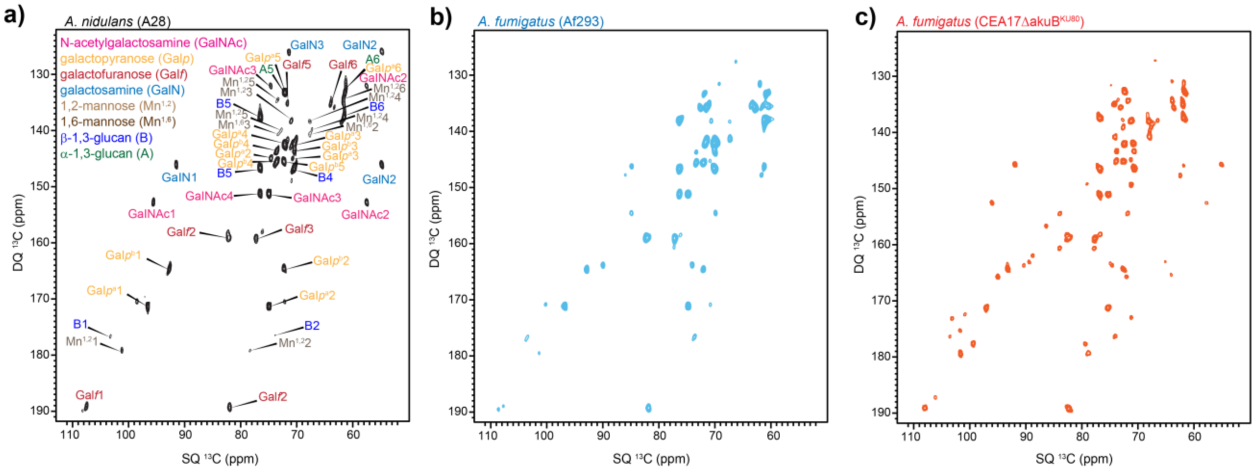


**Supplementary Figure 2. Change is the mobile carbohydrates in different *Aspergillus* fungi.** (**a**) The 2D $^{13}$C-DP INADEQUATE spectra showing all the mobile glucans. (**b**) The mobile spectra representing Af293 strains of *A. fumigatus* showing no $^{13}$C bond for GalNAc, (C1- C2) and GalN, (C1-C2). (**c**) The mobile spectra representing for CEA17ΔakuB$^{KU80}$ with the signals for GalNAc(C1-C2) and GalN(C1-C2). The spectra of A28 and Af293 were measured at 15 kHz and CEA17ΔakuB$^{KU80}$ was measured at 12 kHz.

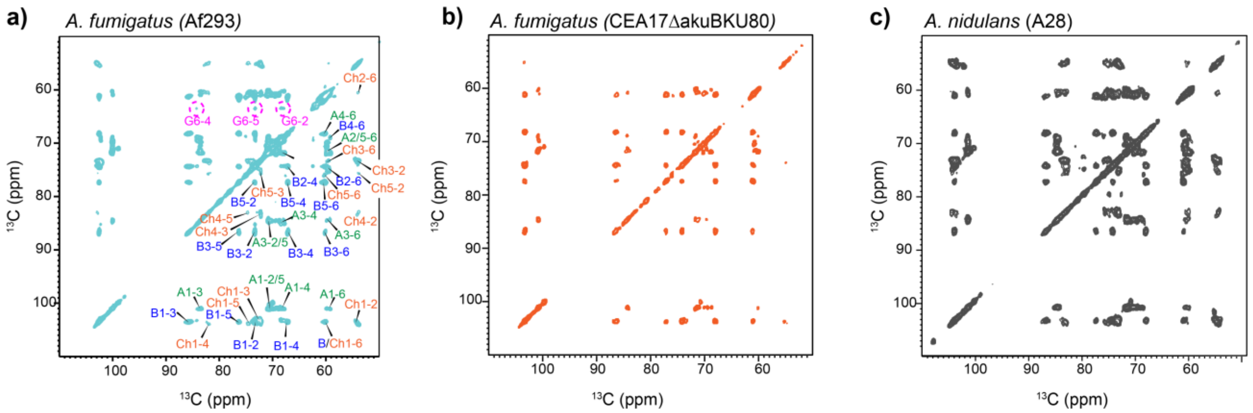


**Supplementary Figure 3. β-1,3/1,4-glucose present in Af293**. (**a**) 2D $^{13}C$-$^{13}C$ CORD spectra showing the rigid components in *A. fumigatus* (Af293) (turquoise) and (**b**) 2D $^{13}C$-$^{13}C$ CORD spectra showing the rigid components for *A. fumigatus* (CEA17ΔakuB$^{KU80}$) in orange spectra. (**c**) 2D $^{13}C$-$^{13}C$ CORD spectra showing the rigid components for *A. nidulans* (A28) in gray spectra. All measurements were obtained using 800 MHz NMR spectrometer.

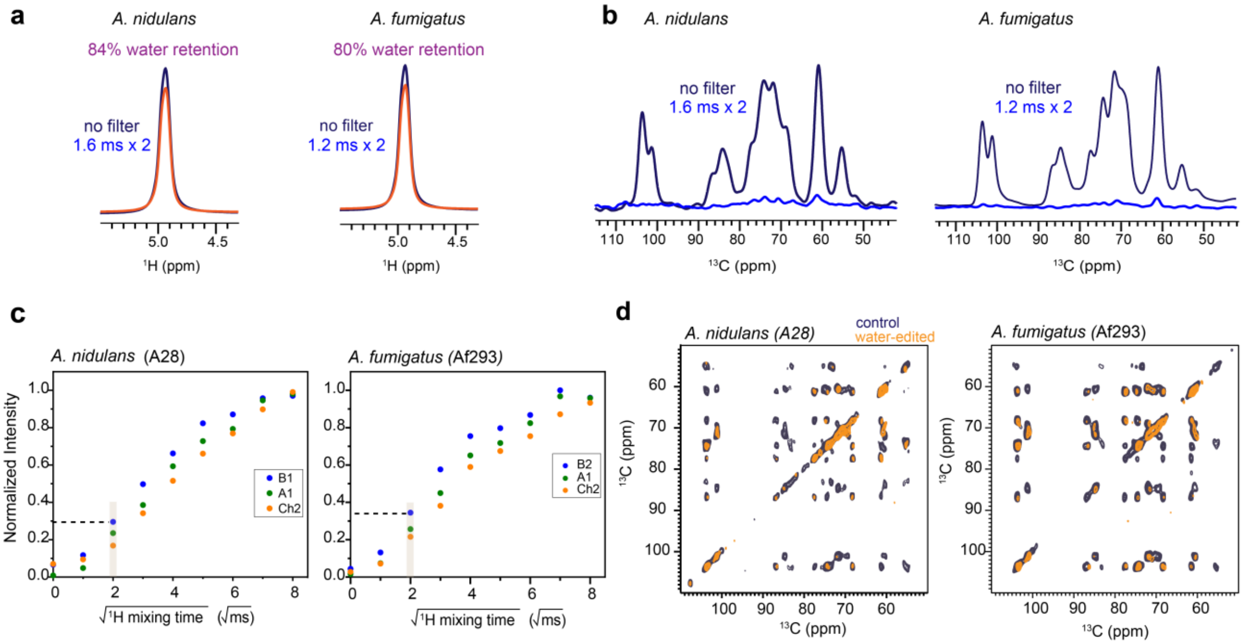


**Supplementary Figure 4. Water-edited analysis for *Aspergillus* cell wall.** (**a**) Proton spectra illustrating the retention of 84% of water magnetization in *A. nidulans* (A28) and 80% in *A. fumigatus* (Af293) after the $^{1}$H-$T_2$ filter. (**b**) Overlay of the $^{13}$C CP spectra where $T_2$ is 1.6 ms × 2 and $T_2$ is 1.2 ms × 2 with no spin diffusion presented in *A. nidulans* (A28) and *A. fumigatus* (Af293) respectively. The $^{1}$H-$T_2$ filtered resulted in a loss of 90% and 89% of the water magnetization in *A. nidulans* (A28) and *A. fumigatus* (Af293), respectively. (**c**) The water to build up curve illustrated for each glucan type which are color coded blue, green and orange for β-1,3-glucan, α-1,3-glucan and chitin respectively. (**d**) Water-edited intensities of carbohydrate carbon sites in *A. nidulans*, A28 (left) and *A. fumigatus*, Af293 (right). All the measurements were taken on a 400 MHz spectrometer at 10 kHz MAS and 280 K.

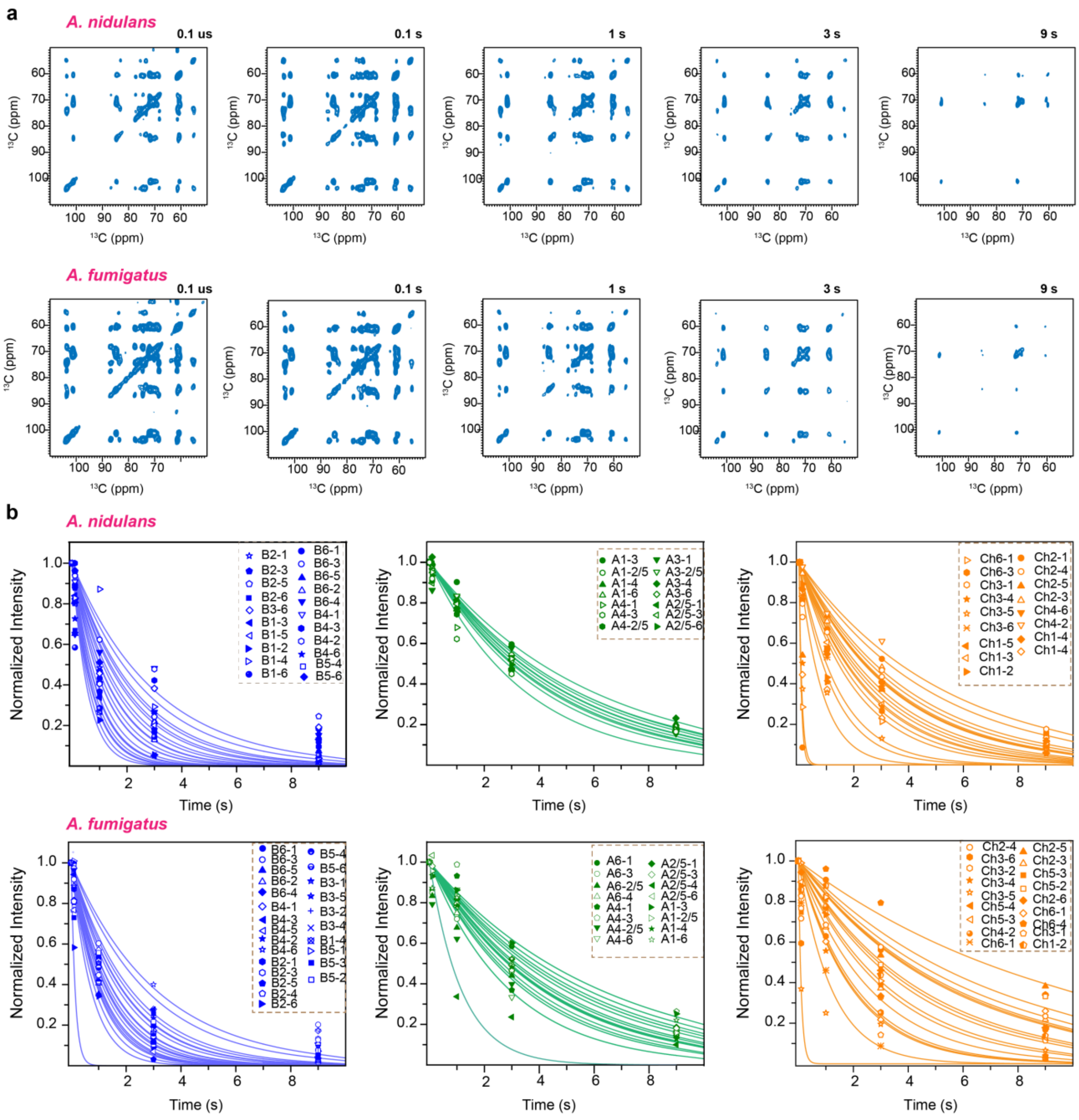


**Supplementary Figure 5. 2D $^{13}$C-$^{13}$C spectra for measuring $^{13}$C-$T_1$ relaxation. a,** The 2D $^{13}$C-$^{13}$C $T_1$ relaxation spectra obtained for examining cellular mobility in *A. nidulans* (top) and *A. fumigatus* (bottom). The spectra are organized in a sequence showing the varying z-filter duration durations; 0 s, 0.1 s, 1 s, 3 s, and 9 s. **b,** $^{13}$C-$T_1$ relaxation curves of *Aspergillus* cell wall polysaccharides**.** The 2D $^{13}$C-$T_1$ relaxation curves are displayed for *A. nidulans* (top) and *A. fumigatus* (bottom). The data are acquired on a 400 MHz (9.4 Tesla) spectrometer with a 10 kHz MAS and at 298 K. The best fit is determined using a single exponential equation. β-1,3-glucan are represented by the blue curves while the α-1,3-glucan and chitin are represented as green and orange curves respectively. Symbols are used for assigning different carbons in the polysaccharides.

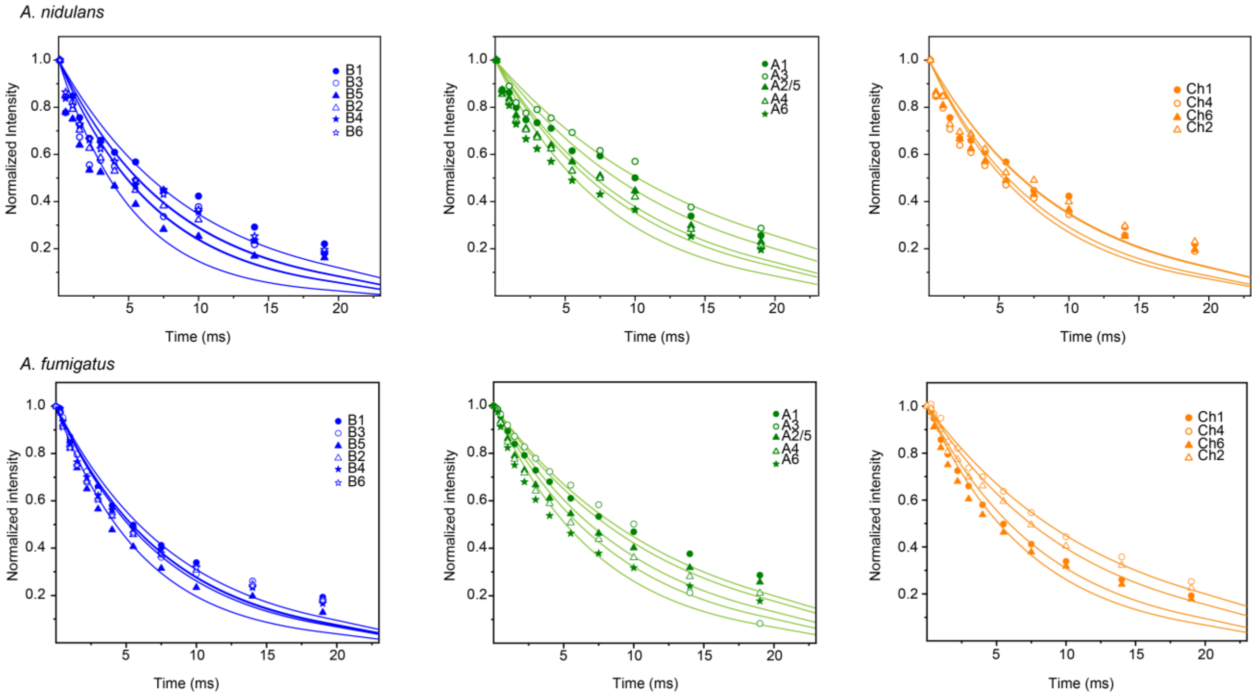


**Supplementary Figure 6. $^1$H-T$_{1\rho}$ relaxation curves of *Aspergillus* cell wall polysaccharides.** The $^1$H-T$_{1\rho}$ relaxation curves are displayed for *A. nidulans* (top) and *A. fumigatus* (bottom). The data are acquired on a 400 MHz (9.4 Tesla) spectrometer with a 10 kHz MAS. The best fit is determined using a single exponential equation. β-1,3-glucan are represented by the blue curves while the α-1,3-glucan and chitin are represented as green and orange curves respectively. Symbols are used for assigning different carbons in the polysaccharides.

**Supplementary Table 1: Salt Solution concentration and minimal media components for *A. nidulans* (A28) and *A. fumigatus* (Af293) culture.**

| Minimal Media | |
|---|---|
| Salt components | g/L |
| $NaNO_3$ | 120.0 |
| KCl | 10.4 |
| $MgSO_4.7H_2O$ | 10.4 |
| $KH_2PO_4$ | 120 |
| $K_2HPO_4$ | 20.9 |
| Glucose | 10.0 |
| Trace element solution | 1ml |

| Trace Elements | g/100ml |
|---|---|
| $CoCl_2.6H_2O$ g | 0.16 |
| $CuSO_4 \cdot 5H_2O$ | 0.16 |
| MnCl2 ·4H2O | 0.5 |
| $(NH_4)6Mo_7O_{24}.4H_2O$ | 0.11 |
| $Na_2EDTA \cdot 4H_2O$ | 0.11 |
| ZnSO4 ·7H2O | 2.2 |
| $H_3BO_3$ | 1.1 |
| $FeSO_4.7H_2O$ | 0.5 |

**Supplementary Table 2. The average cell wall thickness of the three similar fungi**. Results are described as the mean and the standard deviation of 10 individual cells with n=100 measurements in each fungal cell sample. Statistical analysis was performed using one-tailed, paired t-test at 95% confidence level ($p < 0.05$).

| Sample | *A. nidulans* A28 | *A. fumigatus* Af293 | *A. fumigatus* CEA17Δ*akuB*$^{KU80}$ |
|---|---|---|---|
| Average cell wall thickness (nm) | 191 ±16 | 180 ± 23 | 206 ± 54 |

**Supplementary Table 3. Solid- state NMR experimental parameters for *A. nidulans and A. fumigatus*** The experimental parameters include the [1]H Larmor frequency, total experiment time (t), recycle delay (d1), number of scans (NS), The number of points for the direct (td2) and indirect (td1) dimensions, the acquisition time of the direct dimension (aq2) and the evolution time of indirect dimension (aq1), mixing time ($t_m$), and T filter times ($t_z$). * Indicates the water-polysaccharide spin diffusion and the DARR mixing time.

| | Samples | Experiments | (Temp K) | $B_0$ (T) | $\nu_{MAS}$ (kHz) | d1 (s) | NS | td2 | td1 | aq2 (ms) | aq1 (ms) | $t_m$ (ms) | $t_z$ |
|---|---|---|---|---|---|---|---|---|---|---|---|---|---|
| 1D | A28 Af293 | CP | 298 | 18.8 | 15 | 2 | 1024 | 3600 | | 18 | | | |
| | A28 | DP | 298 | 18.8 | 15 | 2 | 512 | 3600 | | 18 | | | |
| | | | | | | 35 | | | | | | | |
| | Af293 | | 293 | | 13.5 | 2 | 128 | 4096 | | 29 | | | |
| | | | | | | 30 | 32 | | | | | | |
| | A28 Af293 | Water-edited | 280 | 9.4 | 10 | 2 | 512 | 2000 | | 16 | | 0, 1, 4,9,16,25,36,49,64, 81,100 | |
| | A28 Af293 | $^{1}H\ T_{1\rho}$ relaxation | 298 | 9.4 | 10 | 2 | 512 | 1400 | | 16 | | | SL (0.1-19 ms) |
| | A28 Af293 | INEPT | 298 | 18.8 | 15 | 3.5 | 1024 | 3200 | | 16 | | | |
| | | | 293 | | 13.5 | 3.0 | 64 | 4400 | | 30.8 | | | |
| 2D | A28 Af293 | 2D CORD | 298 | 18.8 | 15 | 2 | 32 | 3200 | 400 | 16 | 5.2 | 50 | |
| | A28 Af293 | CP *J* INADEQUATE | 298 | 18.8 | 15 | 2 | 32 | 2800 | 472 | 14 | 5.2 | | |
| | A28 A293 | DP *J* INADEQUATE | 298 | 18.8 | 15 | 2 | 32 | 2800 | 472 | 14 | 5.2 | | |
| | A28 Af293 | Pseudo 3D $^{13}C$-$T_1$ | 298 | 9.4 | 10 | 2 | 32 | 2000 | 100 | 16 | 5.5 | 50 | ($10^{-7}$, 0.1, 1, 3, 9) s |
| | A28 Af293 | Water-edited control | 280 | 9.4 | 10 | 2 | 64 | 2000 | 1 | 16 | 5.5 | 0/50* | $T_2$ $10^{-4}$ ms |
| | A28 Af293 | Water-edited | 280 | 9.4 | 10 | 2 | 64 | 2000 | 1 | 16 | 5.5 | 4/50* | $T_2$ 1.6 ms $T_2$ 1.2 ms |
| DNP | A28 | DNP CP | 92 | 14.1 | 10 | 2 | 8 | 2048 | 1 | 10.24 | | | |
| | | N(CA)CX | | 14.1 | 10 | 3.64 | 32 | 1700 | 50 | 12.69 | 3.13 | 100 | |
| | | | | 14.1 | 10 | 3.64 | 32 | 1700 | 50 | 12.69 | 3.13 | 3000 | |
| | | PAR | | 14.1 | 10 | 3.64 | 16 | 1800 | 400 | 13.44 | 7 | 2 | |
| | | | | 14.1 | 10 | 3.64 | 16 | 1800 | 400 | 13.44 | 7 | 20 | |

**Supplementary Table 4. Experimental parameters used for proton detection experiments.** All experiments were performed on 800 MHz (18.8 T) spectrometer and with the MAS frequency of 40 kHz.

| Expt 2D | Samples | Temperature (K) | CP duration (µs) | | D1 | NS | Td2 | Td1 | Aq2 (ms) | aq2 (ms) | Decoupling power | Water suppression (ms) | *J*-evolution (ms) |
|---|---|---|---|---|---|---|---|---|---|---|---|---|---|
| | | | $t_{cp1}$ | $t_{cp2}$ | | | | | | | | | |
| hCH | A28<br>Af293<br>CEA17Δ*akuB*$^{KU80}$ | 304<br>291<br>293 | 200 | 50 | 3 | 8 | 1204 | 512 | 19.98 | 6.4 | slpTPPM [1]<br>(rf 10 kHz) | MISSISSIPI [2]<br>(total duration)<br>200 | ---- |
| refINEPT-HSQC | A28<br>Af293<br>CEA17Δ*akuB*$^{KU80}$ | 304<br>291<br>293 | - | - | 3 | 16 | 2408 | 768 | 39.9 | 9.6 | WALTZ-16 [3] | 200 | 2 |

**Supplementary Table 5. $^{13}C$ and $^{15}N$ chemical shift (ppm) of biomolecules in the cell wall in *A. nidulans* (A28).** Underline represents the $^{13}C$ connectivity with ambiguity and the (-) denotes unidentified.

| Biomolecule | | C1 | C2 | C3 | C4 | C5 | C6 | N | Experimental Methods | Cell Wall Portion | References |
|---|---|---|---|---|---|---|---|---|---|---|---|
| β-1,3-glucan | | 103.6 | 74.3 | 86.6 | 68.1 | 77.4 | 61.1 | | $^{13}C$-$^{13}C$ CORD | Rigid | Shim et al. 2007[4] |
| α-1,3-glucan (A[a]) | | 101.1 | 71.8 | 84.5 | 69.8 | 71.7 | 60.2 | | | | Bhanja et al 2014[5] |
| α-1,3-glucan (A[b]) | | 100.0 | 71.6 | - | - | 71.6 | - | | | | |
| Chitin | a | 104.0 | 55.2 | 73.5 | 83.1 | 75.9 | 60.7 | 126.2 | CORD-$^{13}C$-$^{13}C$ $^{15}N$-$^{13}C$ N(CA)CX | | Fontaine et al. 2011[6] |
| | b | 103.8 | 55.6 | 73.7 | 83.4 | 76.1 | 61.4 | 123.3 | | | |
| | c | 103.5 | 54.5 | 73.6 | 83.0 | 75.4 | 60.9 | 125.5 | | | |
| | d | 103.6 | 55.9 | 73.1 | 82.1 | <u>76.1</u> | 61.6 | | | | |
| Mn[1,2] | | 101.1 | 78.2 | 71 | 67.4 | 73.5 | 61.4 | | $^{13}C$-$^{13}C$ *J*- DP INADEQUATE | Mobile | Latge et al. 1994[7] |
| Mn[1,6] | | 102.5 | 70.8 | 73.3 | 67.4 | 73.3 | 66.2 | | | | Chakraborty et al. 2021[8] |
| Gal*f* | | 107.4 | 81.7 | 77.1 | 82.7 | 71.8 | 63.6 | | | | |
| Gal[a]*p* | | 92.6 | 72.1 | 70.2 | 73.3 | 72.1 | 61.2 | | | | |
| Gal[b]*p* | | 96.7 | 75.0 | 76.6 | 70.4 | 72.7 | 61.6 | | | | Poulhazan et al. 2021[9] |
| GalN | | 91.5 | 54.6 | 71.6 | 81.8 | - | - | | | | Fontaine et al. 2011[6, 10] |
| GalNAc | | 95.6 | 57.3 | 75.2 | 76.5 | - | - | | | | |

**Supplementary Table 6. $^{13}C$ chemical shifts (ppm) of biomolecules in the cell wall of *A. fumigatus* (Af293).** Underline represents the $^{13}C$ connectivity with ambiguity and the (-) denotes unidentified.

| Biomolecule | | C1 | C2 | C3 | C4 | C5 | C6 | Experimental Methods | Cell Wall Portion | References |
|---|---|---|---|---|---|---|---|---|---|---|
| β-1,3-glucan | | 103.6 | 74.3 | 86.6 | 68.1 | 77.3 | 61.1 | $^{13}C$-$^{13}C$ CORD | Rigid (Fontaine et al. 2011[6]) | Shim et al. 2007[4] |
| β-1,4-glucan | | 103.2 | 69.4 | 72.2 | 85.3 | 74.3 | 63.3 | | | Kang et al. 2018[11] |
| α-1,3-glucan ($A^a$) | | 101.1 | 71.8 | 84.5 | 69.8 | 71.7 | 60.2 | | | Bhanja et al 2014[5] |
| α-1,3-glucan ($A^b$) | | 100.0 | 71.5 | - | - | 71.5 | - | | | |
| Chitin | a | 104.0 | 55.2 | 73.5 | 83.1 | 75.9 | 60.7 | CORD-$^{13}C$-$^{13}C$ | | Fontaine et al. 2011[4] |
| | b | 103.8 | 55.6 | 73.7 | 83.4 | 76.1 | 61.4 | | | |
| Mn$^{1,2}$ | | 101.1 | 78.2 | 71 | 67.4 | 73.5 | 61.4 | $^{13}C$-$^{13}C$ *J*- DP INADEQUATE | Mobile | Latge et al. 1994[7] |
| Mn$^{1,6}$ | | 102.5 | 70.8 | 73.3 | 67.4 | 73.3 | 66.2 | | | Latge et al. 1994[7] Chakraborty et al. 2021[8] |
| Gal*f* | | 107.4 | 81.7 | 77.1 | 82.7 | 71.8 | 63.6 | | | |
| Gal$^a$*p* | | 92.6 | 72.1 | 70.2 | 73.3 | 72.1 | 61.2 | | | |
| Gal$^b$*p* | | 96.7 | 75.0 | 76.6 | 70.4 | 72.7 | 61.6 | | | Poulhazan et al. 2021[9] |
| GalN | | 91.5 | 54.4 | 71.6 | 81.8 | - | - | | | Fontaine et al. 2011[6] |
| GalNAc | | 95.4 | 57.2 | 75.2 | 76.5 | - | - | | | |

**Supplementary Table 7. Relative molar composition of *Aspergillus* cell wall.** Molar composition of rigid and mobile components were calculated using the integrals of well-resolved cross peaks of β-1,3-glucan, α-1,3-glucan, and chitin in 2D $^{13}C$-$^{13}C$ CORD spectra and 2D $^{13}C$-$^{13}C$ *J*-INADEQUATE. The results were already normalized by number of scans. (-) is the notation for not detected.

| | **Polysaccharide** | | A28 | | | A293 | | | CEA17Δ*akuB*$^{KU80}$ | | |
|---|---|---|---|---|---|---|---|---|---|---|---|
| | | | Cross-peaks | % | Error % | Cross-peaks | % | Error % | Cross-peaks | % | Error % |
| Rigid Composition | β -1,3-glucan | | B1-3, B1-5, B1-2, B1-4, B2-4, B2-6, B3-5, B3-2, B3-4, B3-6, B5-2, B5-4, B5-6 | 25.2 | 0.1 | B1-3, B1-4, B1-2, B1-6, B1-5, B3-5, B3-2, B3-6, B5-6, B5-4, B2-4, B4-6 | 31.6 | 7.2 | B1-3, B1-4, B1-5, B3-4, B3-5, B3-2, B5-4, B5-2, B2-4 | 50% | 6% |
| | β -1,4-glucose | | G1-4, G1-2, G1-6 | / | / | G1-6, B3-6, G4-5 | 5.2 | 1.6 | / | | |
| | α-1,3-glucan | | A1-2/5, A1-3, A1-4, A3-4, A3-2/5, A2/5-6, A2/5-4 | 44 | 0.9 | A1-4. A1-2/5, A1-4, A1-6, A3-2/5, A2/5-6 | 40.4 | 10.5 | A1-4. A1-2/5, A3-2/5, A3-4 | 42% | 7% |
| | Chitin | | Ch1-4, Ch1-5, Ch1-2, Ch3-2, Ch4-2, Ch4-3, Ch4-5, Ch5-2, Ch5-3, Ch3-2 | 31.2 | 0.1 | Ch1-4. Ch1-2, Ch4-6, Ch4-5, Ch4-3, Ch2-4, Ch2-6 | 22.4 | 4.8 | Ch1-4, Ch1-2, Ch1-5, Ch1-3, Ch4-2, Ch4-2, Ch4-3, Ch4-5, Ch5-3, C3-2, Ch5-2 | 8% | 3% |
| composition | GM | Gal*f* | Gal*f* 1-2, Gal*f* 2-3, Gal*f* 5-6 | 13 | 3 | Gal*f* 1-2, Gal*f* 2-3, Gal*f* 4-5, Galf 3-4, Gal*f* 5-6 | 24.8 | 6.7 | Gal*f* 1-2, Gal*f* 3-4 | 20% | 2% |
| | | $Mn^{1,2}$ | $Mn^{1,2}$1-2, $Mn^{1,2}$2-3, $Mn^{1,2}$ 3-4, $Mn^{1,2}$ 5-6, | 7 | 2 | $Mn^{1,2}$1-2, $Mn^{1,2}$ 3-4, $Mn^{1,2}$ 5-6 | 10.9 | 2.5 | $Mn^{1,2}$1-2, $Mn^{1,2}$4-5 | 24% | 1% |
| | | $Mn^{1,6}$ | $Mn^{1,6}$ 3-4, $Mn^{1,6}$5-6, | 8 | 2 | $Mn^{1,6}$1-2, $Mn^{1,6}$ 3-4, $Mn^{1,6}$2-3 | 20.4 | 6.8 | $Mn^{1,6}$ 1-2, $Mn^{1,6}$ 3-4, | 5.1% | 0.3% |
| | GAG | Gal*p* | Gal1-2, Gal3-4, Gal5-6, Gal2-3 | 29.7 | 9 | Gal1-2, Gal3-4, Gal5-6 | 20.2 | 6.0 | Gal1-2, Gal3-4 | 27% | 2% |
| | | GalNAc | GalNAc 1-2, GalNAc 3-4, GalNAc 2-3 | 20 | 10 | GalNAc 1-2, GalNAc 3-4 | 11.3 | 5.2 | GalNAc 1-2, GalNAc 3-4 | 13% | 3% |
| | | GalN | GalN1-2, GalN3-2 | 16.3 | 5 | GalN | 1.9 | 4.1 | GalN 1-2, GalN3-4 | 6% | 1% |
| other | B1-2 | | | 4 | 1 | B-2, B3-6 | 9.0 | 0.4 | A1-2, A4-5 | 0.83% | 0.1% |
| | A2/5-6 | | | 2 | 1 | A2/5-4 | 1.5 | 0.3 | B1-2, B4-5 | 4% | 1% |

**Supplementary Table 8. Intermolecular interaction of polysaccharides in *A. nidulans* cell wall.** The chemical shifts for the two dimensions of the spectra ($\omega_1$ and $\omega_2$), The cross-peak assignments and the spectral type are documented in the table below.

| Cross Peak | Chemical Shift ($\omega_1$) | Chemical Shift ($\omega_2$) | 2 ms PAR | 20 ms PAR | NCACX (3 s PDSD) |
|---|---|---|---|---|---|
| ChMe-Me' | 23.3 | 22.2 | | x | |
| ChMe'-Me | 21.5 | 22.2 | | x | |
| ChMe-Ch4 | 82.3 | 22.4 | | x | |
| Ch1-ChO | 103.3 | 174.0 | | x | |
| A2-Ch4 | 83.0 | 71.1 | | x | |
| A5-Ch4 | 83.0 | 71.1 | | x | |
| A4-Ch2 | 60.0 | 83.4 | | x | |
| A1-Ch1 | 103.6 | 101.1 | | x | |
| A1-Ch2 | 55.0 | 101.1 | | x | |
| A1-Ch4 | 82.3 | 101.4 | x | x | |
| A1-Ch6 | 60.1 | 100.1 | | x | |
| A2-Ch4 | 71.2 | 83.0 | | x | |
| A5-Ch4 | 71.2 | 83.0 | | x | |
| A6-ChMe | 60.2 | 22.3 | | x | |
| A2-ChCO | 72.1 | 176.8 | | x | |
| A5-ChCO | 72.1 | 176.8 | | x | |
| A3-ChMe | 84.0 | 22.6 | | x | |
| A1-ChMe | 101.4 | 22.3 | | x | |
| Ch2-A1 | 101.3 | 55.8 | | x | |
| Ch2-A6 | 60.0 | 54.5 | | x | |
| Ch3-A1 | 73.3 | 100.1 | | x | |
| Ch4-A2 | 83.3 | 70.2 | x | x | |
| ChMe-A1 | 101.4 | 23.4 | | x | |
| ChMe-A3 | 84.0 | 22.4 | | x | |
| CHMe-A2 | 82.3 | 22.6 | | x | |
| CHMe-A5 | 71.6 | 22.6 | | x | |
| ChN-A3 | 84.8 | 126.5 | | | x |
| A6-B6 | 62..2 | 60.0 | x | x | |
| A1-B2 | 74.4 | 101.3 | | x | |
| A1-B4 | 68.3 | 101.1 | x | x | |
| A1-B6 | 68.8 | 100 | | x | |
| B5-A6 | 60.07 | 77.2 | | x | |
| B6-A1 | 100.1 | 61.2 | | x | |
| B6-A6 | 62.2 | 60.0 | | x | |
| B1-ChMe | 103.6 | 22.4 | | | |
| B6-ChMe | 61.7 | 22.3 | | x | |
| B5-Ch6 | 60.7 | 77.2 | | x | |
| Ch2-B6a | 61.8 | 55.0 | | x | |
| Ch4-B4 | 68.6 | 83.2 | | x | |
| ChN-B3 | 86.1 | 126.1 | | | x |
| ChN-B5 | 77.5 | 126.4 | | | x |
| B5-ChMe | 77.2 | 22.5 | | x | |
| ChMe-B5 | 22.4 | 77.7 | | x | |
| ChMe-B6 | 61.6 | 21.5 | | x | |
| B2-ChCO | 74.8 | 177.1 | | x | |
| B1-ChCO | 103.3 | 177.2 | | x | |

**Supplementary Table 9: Water-edited intensities of the polysaccharides**. The intensity ratios are obtained by comparing the peak intensities between water-edited and the control 2D spectra of the *Aspergillus* cell walls. The error bars represent propagated standard deviations from NMR signal-to-noise ratios, with an error margin typically below 10%,

| β-1,3-glucan | | A28 | Af293 | α-1,3-glucan | | A28 | Af293 | Chitin | | A28 | Af293 |
|---|---|---|---|---|---|---|---|---|---|---|---|
| **103 line** | **103.6** | 0.25±0.03 | 0.31±0.02 | **101 line** | **101.1** | 0.22±0.04 | 0.28±0.03 | **103 line** | **103.4** | 0.26±0.03 | 0.32±0.03 |
| | **86.7** | 0.4±0.1 | 0.40±0.09 | | **84.5** | 0.3±0.1 | 0.20±0.07 | | **83.1** | - | 0.19±0.07 |
| | **77.4** | 0.3±0.1 | 0.5±0.1 | | **71.6** | 0.19±0.06 | 0.20±0.04 | | **61.1** | 0.20±0.09 | 0.32±0.08 |
| | **74.3** | 0.22±0.05 | 0.32±0.03 | | **69.6** | 0.3±0.1 | 0.22±0.06 | | **55.3** | | 0.05±0.02 |
| | **68.1** | 0.3±0.1 | 0.37±0.06 | | **60.7** | 0.30±0.07 | 0.28±0.09 | | | | |
| | **61.1** | 0.19±0.09 | 0.32±0.08 | | | | | | | | |
| **86 line** | **103.5** | 0.3±0.1 | 0.23±0.06 | **84 line** | **101.1** | 0.14±0.08 | 0.18±0.05 | **83 line** | **103.6** | 0.23±0.03 | 0.28±0.08 |
| | **86.6** | 0.4±0.1 | 0.21±0.08 | | **84.5** | 0.22±0.05 | 0.25±0.05 | | **83.7** | - | 0.3±0.1 |
| | **77.3** | 0.40±0.07 | 0.3±0.1 | | **71.6** | 0.20±0.06 | 0.24±0.04 | | **73.4** | - | - |
| | **74.3** | 0.24±0.09 | 0.25±0.07 | | **70.2** | 0.24±0.08 | 0.18±0.06 | | **60.3** | - | 0.26±0.08 |
| | **68.2** | 0.4±0.1 | 0.24±0.07 | | **60.6** | 0.3±0.1 | 0.14±0.06 | | **54.7** | - | 0.25±0.02 |
| | **61.1** | 0.3±0.1 | 0.31±0.09 | | | | | | | | |
| **77 line** | **103.5** | 0.4±0.1 | 0.22±0.07 | **70 line** | **100.9** | 0.31±0.07 | 0.27±0.05 | **75 line** | **103. 6** | 0.23±0.08 | 0.23±0.09 |
| | **86.71** | 0.4±0.1 | 0.16±0.09 | | **84.5** | 0.20±0.09 | 0.30±0.08 | | **83.2** | 0.20±0.08 | 0.19±0.06 |
| | **77.1** | 0.29±0.04 | 0.34±0.04 | | **70.6** | 0.30±0.03 | 0.27±0.02 | | **75.1** | 0.21±0.05 | 0.18±0.03 |
| | **74.4** | 0.24±0.09 | 0.22±0.06 | | **60.8** | 0.30±0.06 | 0.27±0.05 | | **60.8** | 0.15±0.09 | 0.3±0.1 |
| | **68.1** | 0.28±0.09 | 0.25±0.05 | | | | | | **54.8** | 0.17±0.08 | 0.15±0.03 |
| | **61.2** | 0.33±0.06 | 0.24±0.05 | | | | | | | | |
| **74 line** | **103.5** | 0.24±0.04 | 0.25±0.03 | **71 line** | **101.0** | 0.21±0.06 | 0.18±0.03 | **73 line** | **103.6** | 0.27±0.08 | 0.25±0.03 |
| | **86.6** | 0.3±0.1 | 0.35±0.09 | | **84.6** | 0.24±0.09 | 0.23±0.05 | | **83.2** | - | 0.20±0.07 |
| | **77.2** | 0.32±0.08 | 0.07±0.02 | | **71.4** | 0.27±0.03 | 0.22±0.02 | | **60.9** | 0.20±0.08 | 0.18±0.06 |
| | **74.2** | 0.25±0.02 | 0.27±0.02 | | **60.7** | 0.26±0.07 | 0.19±0.03 | | **55.2** | 0.43±0.07 | 0.3±0.1 |
| | **68.2** | 0.32±0.08 | 0.24±0.05 | | | | | | | | |
| | **61.0** | 0.18±0.05 | 0.19±0.05 | | | | | | | | |
| **68 line** | **103.6** | 0.3±0.1 | 0.24±0.05 | **61 line** | **101.1** | 0.19±0.06 | 0.23±0.07 | **61 line** | **103.6** | 0.25±0.09 | 0.16±0.04 |
| | **86.6** | 0.30±0.07 | 0.26±0.08 | | **84** | - | 0.24±0.08 | | **61.0** | 0.26±0.01 | 0.23±0.01 |
| | **77.4** | 0.3±0.1 | 0.32±0.07 | | **71.7** | 0.29±0.06 | 0.21±0.03 | | **55.6** | 0.18±0.08 | - |
| | **74.3** | 0.4±0.1 | 0.22±0.05 | | **61.0** | 0.26±0.02 | 0.23±0.01 | | | | |
| | **68.5** | 0.25±0.06 | 0.20±0.03 | | | | | | | | |
| | **61.1** | 0.29±0.09 | 0.21±0.05 | | | | | | | | |
| **61 line** | **103.6** | 0.25±0.09 | 0.24±0.05 | | | | | **55 line** | **103.8** | 0.13±0.01 | 0.09±0.03 |
| | **86.6** | 0.30±0.09 | 0.4±0.1 | | | | | | **83.2** | - | 0.12±0.03 |
| | **77.5** | 0.32±0.09 | 0.34±0.05 | | | | | | **75.6** | - | 0.4±0.1 |
| | **74.7** | 0.22±0.07 | 0.27±0.05 | | | | | | **73.2** | 0.14±0.06 | 0.3±0.1 |
| | **68.3** | 0.21±0.07 | 0.21±0.03 | | | | | | **60.4** | 0.11±0.05 | 0.20±0.06 |
| | **61.0** | 0.26±0.02 | 0.23±0.01 | | | | | | | | |
| **Average** | | **0.29** | **0.27** | | | **0.24** | **0.23** | | | **0.20** | **0.22** |
| **n** | | **36** | **36** | | | **21** | **22** | | | **17** | **26** |

**Supplementary Table 10. $^1$H-$T_{1\rho}$ relaxation times of polysaccharides.** A single exponential equation was used to fit the $T_1$ data $I(t) = e^{-t/T_I}$. A single exponential equation was used to fit the $T_{1\rho}$ data: $I(t) = e^{-t/T_{I\rho}}$. Error bars are standard deviations of the fit parameters.

| Sample Type | Cross peaks | $^1$H-$T_{1\rho}$ (ms) | Average |
|---|---|---|---|
| **A28** | B1 | 9.4 ± 0.8 | 7.5 ± 0.7 |
| | B3 | 7.0 ± 0.8 | |
| | B5 | 5.2 ± 0.6 | |
| | B2 | 7.0 ± 0.7 | |
| | B4 | 8.0 ± 0.7 | |
| | B6 | 8.1 ± 0.7 | |
| | A1 | 12.5 ± 0.9 | 11.0 ± 0.8 |
| | A3 | 14.7±0.9 | |
| | A2/5 | 10.3 ± 0.8 | |
| | A4 | 9.6 ± 0.9 | |
| | A6 | 8.1 ± 0.7 | |
| | Ch1 | 9.4 ± 0.8 | 8.7 ± 0.8 |
| | Ch3 | 7.6 ± 0.7 | |
| | Ch5 | - | |
| | Ch6 | 8.1 ± 0.7 | |
| | Ch2 | 9.6 ± 0.8 | |
| **Af293** | B1 | 8.5 ± 0.4 | 7.5 ± 0.4 |
| | B3 | 7.8 ± 0.4 | |
| | B5 | 6.1 ± 0.3 | |
| | B2 | 7.4 ± 0.4 | |
| | B4 | 7.8 ± 0.4 | |
| | B6 | 7.4 ± 0.5 | |
| | A1 | 12.5 ± 0.6 | 10.1 ± 0.6 |
| | A3 | 11.6 ± 0.7 | |
| | A2/5 | 10.0 ± 0.7 | |
| | A4 | 8.8 ± 0.6 | |
| | A6 | 7.4 ± 0.5 | |
| | Ch1 | 8.5 ± 0.4 | 9.8 ± 0.5 |
| | Ch4 | 12.5 ± 0.4 | |
| | Ch6 | 7.4 ± 0.5 | |
| | Ch2 | 10.8 ± 0.5 | |

**Supplementary Table 11. 2D $^{13}$C-$T_1$ relaxation times of polysaccharides within *Aspergillus* cell walls.** Data are presented for the *A. nidulans* (A28) and *A. fumigatus* (Af293). The bold values indicate average measurements for each polysaccharide within each sample. The relaxation times were obtained through 2D $^{13}$C-$^{13}$C correlation experiments and are fit using single exponential equation $I(t) = e^{-t/T_I}$. Error bars are standard deviations of the fit parameters.

| β-1,3-glucans cross peaks | *A. nidulans* A28 | *A. fumigatus* Af293 | Chitin Cross peaks | A28 | Af293 | α-1,3-glucans cross peaks | A28 | Af293 |
|---|---|---|---|---|---|---|---|---|
| B6-1 | 1.4±0.3 | 1.4±0.5 | Ch2-1 | - | 5.4±0.5 | A1-3 | 5.8±0.4 | 6.3±0.2 |
| B6-3 | 1.9±0.3 | 1.7±0.2 | Ch2-4 | 2.9±0.9 | 3.1±0.9 | A1-2/5 | 5.1±0.2 | 4.6±0.3 |
| B6-5 | 1.2±0.2 | 1.0±0.2 | Ch2-3 | - | 3.3±0.7 | A1-4 | 4.5±0.4 | 4.2±0.2 |
| B6-2 | 1.2±0.4 | 1.1±0.2 | Ch2-3 | 3.9±0.4 | 3.3±0.7 | A1-6 | 4.4±0.4 | 5.2±0.7 |
| B6-4 | 1.9±0.7 | 1.5±0.4 | Ch4-6 | 2.9±0.5 | - | A3-1 | 5.3±0.7 | - |
| B4-3 | 2.0±0.7 | 2.1±0.2 | Ch4-2 | 4.7±0.6 | - | A3-2/5 | 4.9±0.2 | - |
| B4-2 | | 1.9±0.7 | Ch1-4 | 2.7±0.6 | - | A3-4 | 5.1±0.5 | - |
| B4-6 | 1.6±0.7 | 2.8±0.7 | Ch1-5 | 0.8±0.5 | 2.6±0.4 | A3-6 | 4.4±0.3 | - |
| B2-1 | 1.7±0.1 | 1.7±023 | Ch1-3 | 2.5±0.5 | - | A2/5-1 | 4.8±0.1 | 4.9±0.2 |
| B2-3 | 1.5±0.2 | 1.3±0.2 | Ch1-2 | 2.3±0.3 | 4.2±0.8 | A2/5-3 | 4.6±0.4 | 5.4±0.3 |
| B2-5 | 1.0±0.5 | 0.9±0.2 | Ch6-1 | 1.4±0.3 | 2.1±0.7 | A2/5-6 | 4.4±0.4 | 4.4±0.3 |
| B2-6 | 0.9±0.3 | 0.7±0.3 | Ch3-1 | 2.1±0.5 | 2.0±0.4 | A4-1 | 4.3±0.6 | - |
| B5-4 | 0.8±0.1 | 1.5±0.3 | Ch5-3 | - | 4.4±0.4 | A4-3 | 4.1±0.4 | 5.8±1.4 |
| B5-1 | - | 1.3±0.2 | Ch5-2 | - | 3.7±0.4 | A4-2/5 | 3.4±0.6 | 2.9±0.7 |
| B5-3 | - | 1.7±0.5 | Ch3-4 | 3.7±0.9 | 2.6±0.6 | A6-3 | | 3.8±0.6 |
| B5-6 | 1.3±0.2 | 1.5±0.5 | Ch3-5 | 1.2±0.7 | - | A6-2/5 | | 3.5±0.6 |
| B3-6 | 2.2±0.7 | - | Ch3-6 | 0.1±0.5 | 0.1±0.04 | A6-1 | | 5.2±0.7 |
| B1-3 | 0.9±0.2 | - | Ch3-2 | 2.6±0.6 | 2.0±0.8 | A6-4 | | 4.2±0.4 |
| B1-5 | 0.8±0.2 | - | Ch4-1 | | 4.1±0.6 | A4-6 | | 3.6±0.7 |
| B1-4 | 3±0.7 | 1.3±0.1 | Ch4-3 | | 4.0±0.4 | A2/5-4 | | 1.2±0.3 |
| B1-6 | 0.6±0.3 | - | Ch1-6 | | 1.2±0.2 | | | |
| B1-6 | 0.6±0.3 | - | | | | | | |
| B3-1 | - | 1.2±0.1 | | | | | | |
| B3-5 | - | 1.6±0.3 | | | | | | |
| **Average** | **1.39** | **1.48** | | **2.41** | **3.0** | | **4.65** | **4.35** |
| n | 19 | 19 | | 14 | 16 | | 14 | 15 |

**Supplementary Table 12. $^{1}$H and $^{13}$C chemical shifts of *A. nidulans* polysaccharides.** For each carbon site, the $^{13}$C and $^{1}$H chemical shifts are shown in the top and bottom rows, respectively. All $^{1}$H proton chemical shifts were following Safeer et al. 2023[12]. Chemical shifts of NRa, NRb, Rb, B, and A are following Ehren et al. 2020[13]. Chemical shifts of Mn, Galf, Galp, GalN, GalNAc are following values from Latgé et al. 1994[7]and Chakraborthy et al. 2021[8].

| Biomolecule | C1 | C2 | C3 | C4 | C5 | C6 | C8 |
|---|---|---|---|---|---|---|---|
| α-1,3-glucan | 101.1<br>5.4 | 71.8<br>3.7 | 84.5<br>3.6 | 69.8<br>3.7 | 71.7<br>3.8 | 60.9<br>3.8 | --- |
| β-1,3-glucan | 103.6<br>4.9 | 74.3<br>3.6 | 86.7<br>3.6 | 68.1<br>3.4 | 77.3<br>3.4 | 60.9<br>3.8 | --- |
| Chitin | 103.7<br>4.0 | 55<br>3.8 | 73.5<br>3.6 | 83<br>3.6 | 75.6<br>3.6 | 60.9<br>3.8 | 22.8<br>2.1 |
| NR[a] | 98.6<br>5.0 | | | | | | --- |
| R[b] | 98.4<br>4.9 | --- | --- | --- | --- | --- | --- |
| NR[b] | 103.1<br>4.7 | 73.8<br>3.9 | | | | | |
| B | 103.1<br>4.7 | 73.8<br>3.9 | | | | | |
| Mn[1,2] | 101.1<br>5.2 | 78.6<br>4.0 | 71.1<br>3.8 | 67.4<br>3.7 | 73.7<br>4.1 | 61.7<br>3.8 | --- |
| Mn[1,6] | 102.6<br>5.0 | 70.7<br>3.9 | 73.2<br>4.1 | 67.5<br>3.7 | 73.7<br>4.1 | 66.3<br>3.7 | --- |
| Gal*f* | 107.4<br>5.2 | 81.7<br>4.1 | 77.7<br>4.0 | 83.2<br>4.0 | 71.5<br>3.8 | 63.5<br>3.7 | --- |
| Gal[a]*p* | 92.6<br>5.2 | 72.1<br>3.5 | 70.2<br>3.4 | --- | 72.1<br>3.8 | 61.3<br>3.8 | --- |
| Gal[b]*p* | 96.7<br>5.4 | - | 76.5<br>3.4 | 70.3<br>3.4 | - | 61.6<br>3.8 | --- |
| GalN | | 54.4<br>3.2 | | 81.8<br>4.1 | - | - | |
| GalNAc | 96.4/95.6<br>4.6 | 57.2<br>3.7 | --- | 76.9<br>4.0 | - | - | 22.7<br>2.0 |
